# Systematic investigation of synthetic operon designs enables prediction and control of expression levels of multiple proteins

**DOI:** 10.1101/2022.06.10.495604

**Authors:** Daniel Gerngross, Niko Beerenwinkel, Sven Panke

**Author notes:** Centre for Genomic Regulation (CRG), The Barcelona Institute of Science and Technology, Barcelona, Spain.

## Abstract

Controlling the expression levels of multiple recombinant proteins for optimal performance is crucial for synthetic biosystems but remains difficult given the large number of DNA-encoded factors that influence the process of gene expression from transcription to translation. In bacterial hosts, biosystems can be economically encoded as operons, but the sequence requirements for exact tuning of expression levels in an operon remain unclear. Here, we demonstrate the extent and predictability of protein-level variation using diverse arrangements of twelve genes to generate 88 synthetic operons with up to seven genes at varying inducer concentrations. The resulting 2772 protein expression measurements allowed the training of a sequence-based machine learning model that explains 83% of the variation in the data with a mean absolute error of 9% relative to reference constructs, making it a useful tool for protein expression prediction. Feature importance analysis indicates that operon length, gene position and gene junction structure are of major importance for protein expression.

## Introduction

Synthetic biology is establishing itself as a technological approach to design biological systems that address challenges in important industrial areas, such as the production of fine chemicals, fuels and bulk chemicals from renewable resources and the improvement of human health based on smart cell-based diagnostic tools and therapies^1,2^. The required molecular basis for this approach consists of multi-protein systems. For instance, a biocatalytic network to produce a molecule of interest can consist of many enzymes^3,4^, or ultimately therapeutic synthetic circuits can contain multiple regulatory proteins to achieve the intended function^5,6^, and many of these proteins can be heterologous.

Enzymes or regulatory proteins in multi-protein systems often display broadly different kinetic properties such as specific rate constants or affinities. In natural systems, the various proteins evolved so that these properties are adjusted to ensure an organism’s survival and proliferation. However, synthetic systems typically include proteins from various sources or proteins whose original properties may have been substantially altered. Additionally, expressing recombinant genes in alternative chassis might expose them to previously unknown influences, such as inhibitions of degradation.

Poor integration of the properties of recombinant proteins in the system or even general lack of knowledge about the decisive system performance parameters often leads to the situation that the performance of synthetic systems at an early prototype stage is typically not even close to desired levels. Improvement can come from combinatorial or semi-rational variations of relative protein levels^7,8^ to improve a specific target property, which can require the analysis of large libraries. This process could be much streamlined if the behavior of library members could be better predicted, which requires tangible control over the protein expression level of each of the system components.

Most commonly, promoters^9^ and ribosome binding sites (RBSs) are modified to vary protein expression levels^7,10^. For current synthetic multiprotein systems, this means that each protein is encoded as an individual transcription unit to allow for tuning individual protein levels. When the protein system becomes larger, the repertoire of constitutive promoters that are sufficiently distinct from each other to prevent homologous recombination and a resulting genetic instability might no longer be large enough^11^. Also, constitutive expression might lead to accelerated loss of function due to its continuous impact on growth. Using multiple inducible promoters, each with its own regulatory protein^12^, can lead to similar repertoire issues and an additional level of complexity (and cost) at the level of induction.

One way to mitigate this issue is to build synthetic operons. Synthetic operons are collections of recombinant genes that can be transcribed into a single mRNA, and therefore allow controlling several genes from one promoter^13^. In fact, operons are frequently applied in synthetic multiprotein systems. They are also a common natural genetic organization principle: in the bacterial model chassis *Escherichia coli* approximately one third of all transcription units encode operons and more than 50 % of all genes are encoded in operons with three or more protein coding sequences^14^. However, organization of genes in operons expands the list of poorly quantifiable factors that influence protein expression.

The design of a synthetic operon often starts with the choice of gene order before one performs further optimization, for instance, via RBS tuning^7^. Initial operon gene orders typically follow ad-hoc rules like the ordering according to appearance of encoded enzymes in a pathway^15,16^, the order in natural operons^7,17^ – if an equivalent exists – or an arbitrary choice^18,19^. Importantly, some of the elements that are known to influence protein expression levels can strongly depend on the position of the gene within the operon. A prominent example is translational coupling, which describes the effect of the translation efficiency of a gene on the translation of the gene immediately downstream^20,21^. Also, RNases like RNase E specifically initiate 5’-end-dependent degradation of downstream RNA depending on specific motifs ^22^, which in turn can vary protein expression as a function of the gene’s position relative to the 5’ end. Additionally, there could be several (unintended) promoters or terminators associated with certain parts of the operon, so positioning this part at different places in an operon will lead to a distribution of concentrations of subsections of the whole operon transcript^23,24^.

Importantly, starting with a gene order without knowing its potential influence on protein expression levels can lead to unintentional constraints on the variability of expression levels when other elements like promoters or RBSs are subsequently varied for expression level finetuning. For example, some genes might be locked in settings that allow only for high or for low expression levels.

Consequently, we argue that ad-hoc operon design choices, such as, for example, of gene order, ignore an important layer of controlling recombinant protein levels. This is probably due to the lack of comprehensive operon models, which, in turn, is the consequence of limited data based on systematic gene order variation of synthetic operons.

Here, we aim to develop a rational approach to the design of multi-protein systems in synthetic biology by implementing a predictive model for protein expression in operons based only on DNA sequence features, which are rapidly accessible via existing bioinformatics tools. The model is derived from a hitherto unseen dataset capturing variation of gene order in synthetic operons. We establish a workflow to generate a large set of synthetic operons and measure differential protein expression of up to seven encoded reporter proteins simultaneously. Our measurements together with sequence-derived features corresponding to each operon design enable the construction of a machine learning model that is able to predict expression levels with sufficient accuracy. Additionally, we assess the importance of the sequence features for prediction. Finally, we provide open access to the full model and data set for future research and improvement as well as an online tool that allows predicting protein expression levels from user-provided operon sequences.

## Results

### Synthetic operon design and construction

We designed the synthetic operons investigated in this study using a LacI repressible T7 RNA polymerase expression system which can be induced with isopropyl β-D-1-thiogalactopyranoside (IPTG) (see **Figure 1A**). To facilitate comparison of the expression of various proteins across different operon designs, different positions, and different inducer concentrations, we report protein expression (PE) as differential protein expression (DPE). We define DPE as the PE resulting from one gene encoded in a multi-gene operon relative to the PE of the same gene (with the same RBS) encoded in a monocistronic transcription unit at an inducer concentration of 400 μM (see **Figure 1B**). For instance, the DPE of protein A in the operon coding for A, C, and B (DPE_A,ACB_) is the absolute PE of A in the operon ACB (PE_A,ACB_) divided by the absolute PE of A in the monocistronic construct (PE_A,A_).

**Figure 1.**
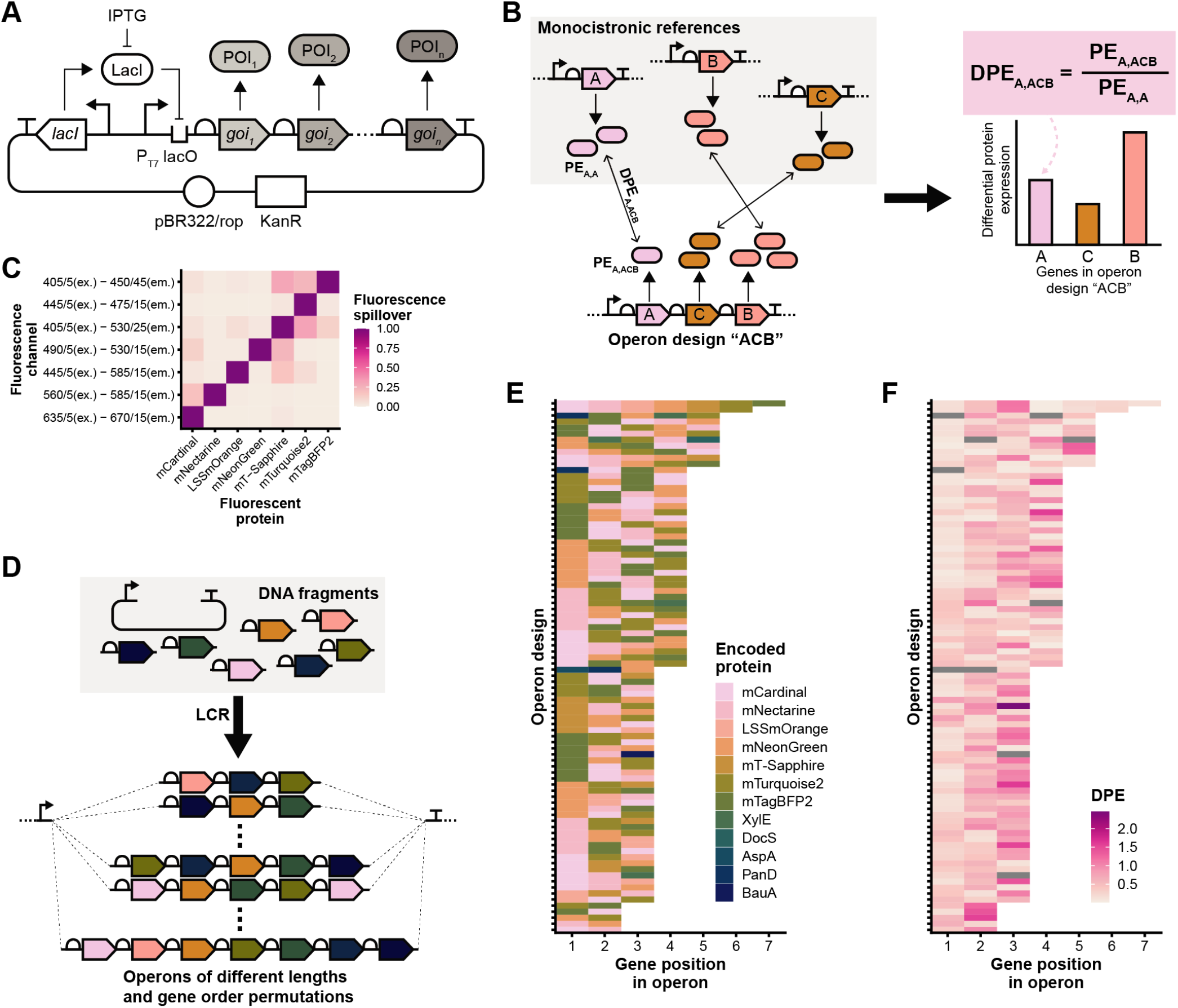
Design, assembly, and measurement of synthetic operons. (**A**) Plasmid design of the operon expression constructs. The backbone is based on the pSEVA291 plasmid containing an operon expression and regulatory cassette. The operon, containing the genes of interest (*goi_1_* to *goi_n_*), is transcribed from a T7 promoter (P_T7_) which is regulated by the LacI inhibitor through interaction at the *lac* operator (*lacO*) and induced by addition of IPTG. The induction leads to the expression of the proteins of interest (POI_1_ to POI_n_). (**B**) For all reported data of the constructed operons, the expression of proteins (PE) encoded in operons was compared to the expression of the same protein encoded in a monocistronic context to obtain the differential protein expression (DPE). (**C**) Spillover matrix of the fluorescent proteins (FPs) used as reporters. The vertical axis indicates the fluorescence channels (excitation and emission wavelengths with respective band width in nm). The horizontal axis indicates the measured FP. The color of each tile indicates the fluorescence intensity in each channel relative to each FP’s target channel. (**D**) Using the ligase cycling reaction (LCR)^35^, plasmid backbone and translation units (ribosome binding site and protein coding sequence) were assembled in various combinations to obtain synthetic operons with various gene order permutations containing up to seven genes. (**E**) Diversity of 88 bi-to heptacistronic operon designs. The colored rectangles represent the genes coding for seven FPs (mCardinal to mTagBFP2) and five non-FPs (XylE to BauA). Each horizontal line of rectangles represents one operon design with a different gene at each position of the operon. (**F**) Differential protein expression for each protein encoded in the operons depicted in D. The color of each rectangle indicates the mean of the DPE across replicates of experiments with an IPTG concentration of 400 μM. Gray rectangles indicate non-FPs that were not measured in the fluorescence-based assays.

For rapid readout of PE for a variety of genes, we used fluorescence of various proteins as a proxy for PE allowing us to determine multiple PEs simultaneously. We used the seven fluorescent proteins (FPs) mCardinal^25^, mNectarine^26^, LSSmOrange^27^, mNeonGreen^28^, mT-Sapphire^29^, mTurquoise2^30^, and mTagBFP2^31^. Using a flow cytometry setup (see Methods), we determined the optimal excitation laser-emission filter combination for each FP resulting in minimal spillover to other fluorescence channels and thus allowing for orthogonal measurements of multiple FPs in a single cell at once (see **Figure 1C**).

We also used five additional genes (non-FPs) of different length encoding for proteins of various functionalities to generate additional diversity in terms of gene and operon length as well as gene distances to the respective operon ends in the set of synthetic operons (encoded proteins: XylE^32^, DocS^33^, AspA^34^, PanD^34^, BauA^34^; see **Figure 1E** and **Figure S2**, Supplementary Information). Coding sequences of all FP genes were chosen to minimize potential homologous stretches of DNA and decrease similarity between the genes within a synthetic operon (see **Figure S2**, Supplementary Information).

An efficient way to generate defined rearrangements in a linear sequence of genetic parts is assembly via the ligase cycling reaction (LCR)^35^. The LCR allows specifying the order of the DNA parts to be assembled by only adding specific bridging oligos to the assembly reaction (**Figure 1D**). Here, each of the DNA parts already contains an RBS and a protein coding sequence. The RBSs were designed using the RBS Calculator^36^ with a target relative translation initiation rate (TIR) of 100,000 in the monocistronic context, resulting effectively in TIRs between approx. 10,000 and 150,000 (see **Figure S2**, Supplementary Information). The synthetic RBSs are approx. 32 bases long and are the only sequence separating stop and start codon of two subsequent protein coding sequences in the operon context. Using the same 12 DNA parts, we constructed 88 operons ranging in length from 2 to 7 genes (see **Table S4**(Supplementary Information)/**Figure 1E**). The backbone was based on the pSEVA291 plasmid^37^ containing a pBR322-ROP origin of replication, a kanamycin resistance gene, the gene for the LacI repressor, and a *lac* operator-controlled T7 promoter upstream of the operon insertion site (see **Figure 1A**). An *E. coli* MG1655 (DE3) strain, which expresses the necessary T7 RNA polymerase from a phage lambda lysogen upon induction with IPTG, was used as expression host.

### Rearrangement of synthetic operon genes leads to diverse protein expression patterns

When investigating PE in the different operons, we found that gene permutations cause large changes in protein expression (**Figure 1F** and **Figure 2B**). DPE varies approximately 600-fold (between 0.004 and 2.5) at induction with 400 μM IPTG, depending on operon design and investigated protein.

**Figure 2.**
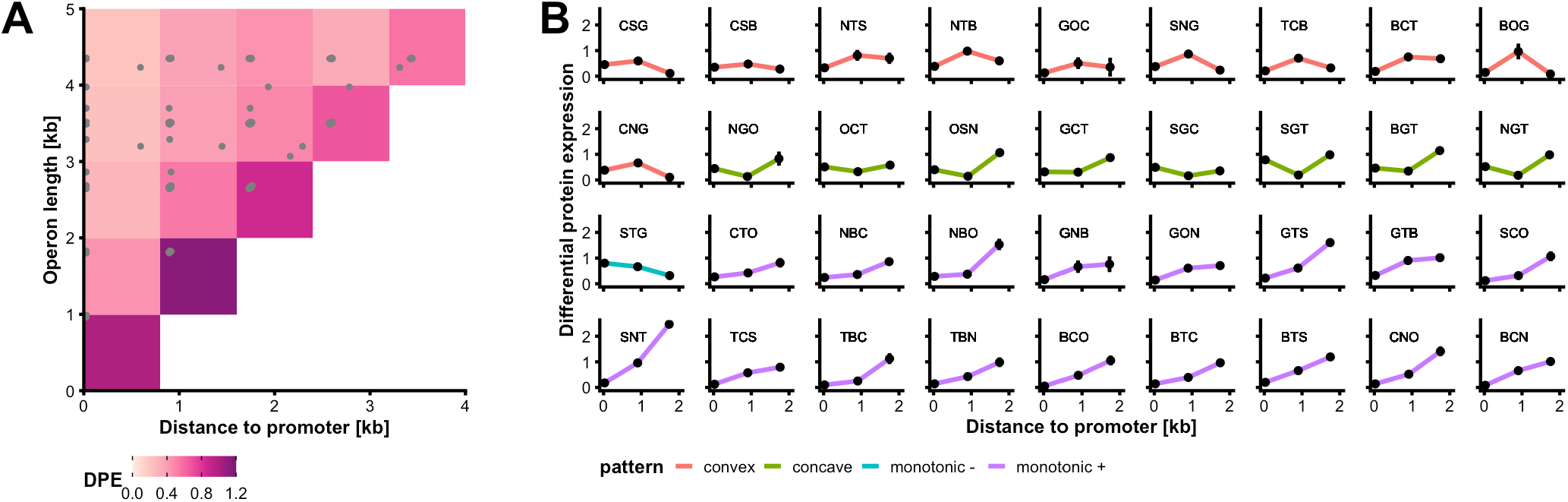
Effect of operon length, gene distance to promoter, and gene order on DPE. (**A**) Summary DPE matrix for operon constructs shorter than 5 kb (86 operons designs) and monocistronic references. The horizontal axis indicates the distance of a gene’s start codon to the transcription start site (TSS) in kb. The vertical axis indicates the operon length between TSS and terminator in kb. The color of each tile indicates the mean DPE value of all encoded proteins whose genes fall within its boundaries. The bottom left tile contains only the monocistronic reference constructs and has therefore per definition a value of 1. The underlying individual operon length and promoter distance values of each operon design are indicated by grey points. (**B**) PE patterns resulting from gene position in all 36 constructed tricistronic operon designs. Each graph represents one operon design indicated by the three letters above each graph (C: mCardinal, N: mNectarine, O: LSSmOrange, G: mNeonGreen, S: mT-Sapphire, T: mTurquoise2, B: mTagBFP2). For instance, the operon design CSG codes for three FPs in following order: mCardinal, mT-Sapphire, mNeonGreen. The vertical axis indicates the DPE. The horizontal axis indicates the respective FP gene’s start codon distance to the TSS in kb. The line connecting the points is meant to illustrate the expression level pattern across each operon. The color of the line represents the four categories of patterns: concave (second position highest expressed; red), convex (second position lowest expressed; green), monotonic decreasing (cyan), and monotonic increasing (purple). The points indicate the mean value of at least three biological replicates for each measurement. The error bars indicate one standard deviation of the mean in each direction. The inducer concentration in all plotted experiments (A and B) was 400 μM IPTG.

Overall, genes encoded in longer operons have lower expression levels. Remarkably, the expression level is higher when genes are located closer to the 3’-end of the operon (see **Figure 2A** and **Figure S3**, Supplementary Information). The data set in **Figure 2A** contains the average differential protein expression of all 86 operon designs with up to 5 FPs. While the mean DPE of genes in these operon variants shows a consistent increase with increasing distance from the promoter in operons of equal length, the individual DPE patterns vary highly between different operon designs. This can be seen for instance in the set of tricistronic operons with varying gene order. This set, containing 36 out of 210 possible tricistronic permutations of the seven FPs, comprises four fundamental DPE patterns: concave, convex, monotonic increasing, and monotonic decreasing (see **Figure 2B**). For longer operons, the observed patterns divert further into complex patterns (see **Figure S3B** for all 88 operon designs, Supplementary Information).

When varying inducer concentrations, the general trend of DPE patterns remains mostly the same (see **Figure S4**, Supplementary Information). In general, the differences in DPE between the genes in the same operon become less pronounced with lower inducer concentrations. However, in some cases, the protein with the highest DPE in a given operon design becomes lower than the DPE of other encoded proteins for lower inducer concentrations (see **Figure S4B**, Supplementary Information).

### Random forest model to predict PE based on synthetic operon sequence

In order to learn more about the most impactful operon design elements and enable predictive design of synthetic operons we used machine learning to construct a model that predicts DPE from the operon sequence. The constructed operon designs contain a set of sequence-dependent elements that were shown previously to potentially influence PE. These features include RBS-dependent translation initiation rate^36^, gene position in the operon^38,39^, mRNA folding energy^40^, structural RNA elements^41,42^, RNase motifs^22,42^, and basic DNA sequence features like GC content and sequence length. A full list of the sequence elements we used as features for our model is available in **Table S5** (Supplementary Information). In view of future use of the model with other genes and new operon designs, all the selected features can be readily extracted from a given sequence using freely available bioinformatics tools also listed in this table.

We used these features, 19 in total, and DPE as response variable to train a random forest regression model. To evaluate model performance, we applied 5-fold cross validation with an 80/20% train/test set split each. Hyperparameters were learned separately within each training set. The set members were sampled according to unique combinations of operon, gene, and induction level to avoid that biological replicates can be simultaneously part of the training and test set (see Methods).

The 5-fold cross-validation resulted in a mean R^2^ of 0.86 (standard deviation of 0.02) and a mean MAE of 0.089 (standard deviation of 0.007) between predicted and measured DPE (**Figure 3A–C**). These results indicate that the estimated model is capable of robustly predicting DPE in synthetic operons. With an MAE below 0.09 on the scale of DPE values (less than 9% relative to the reference PE), we consider the model performance sufficient for the first step in operon design before further fine-tuning is performed. The set of 9 selected operon designs of **Figure 3D** illustrates how closely predictions resemble the measured patterns of increased or decreased DPE along the different positions in each operon (see **Figure S5** for all predicted operon designs, Supplementary Information).

**Figure 3.**
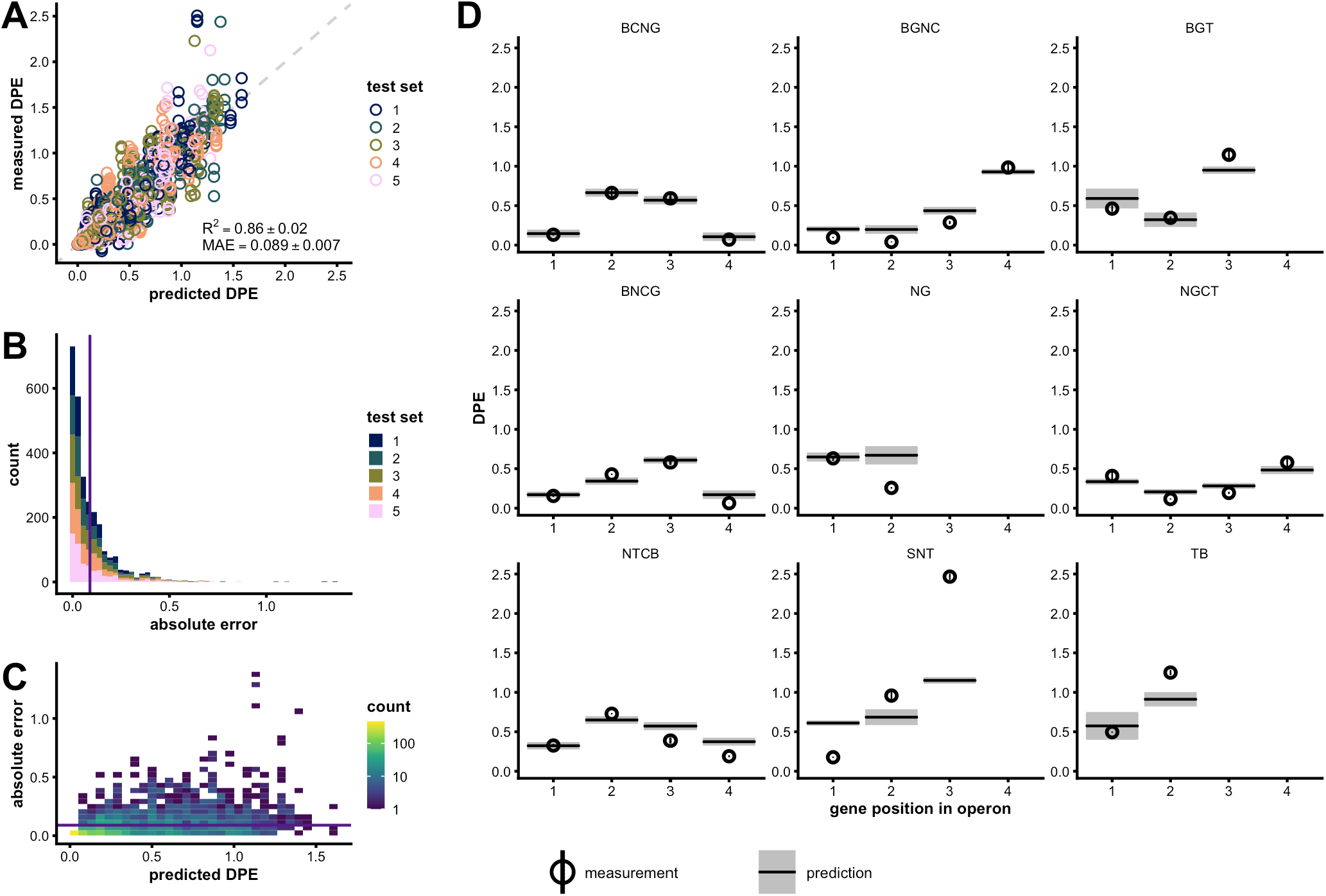
Performance of a random forest regression model in predicting DPE based on sequence features. (**A**) Measured DPE plotted against predicted DPE for each gene using the five test sets of a 5-fold cross-validation of the model. The mean coefficient of determination (R^2^) for five models is 0.86 with a standard deviation of 0.02. The mean of the five mean absolute errors (MAE) is 0.089 with a standard deviation of 0.007. The dashed diagonal line indicates the optimal position of points where a prediction would exactly match the measurement. (**B**) Distribution of absolute errors for the five test sets. The purple line marks the mean MAE. (**C**) Absolute errors compared to corresponding measurements. The colored squares indicate how many values there are within the indicated area of error and DPE values. The purple line marks the mean MAE. (**D**) Individual predictions and measurements for nine randomly chosen operons from the test sets that contain predictions for all their genes (**Figure S5** shows all operons, Supplementary Information). Each panel represents one operon design indicated by the order of the letters above each panel (C: mCardinal, N: mNectarine, O: LSSmOrange, G: mNeonGreen, S: mT-Sapphire, T: mTurquoise2, B: mTagBFP2). The horizontal axes show the position of a gene within an operon. The circles indicate the mean value of at least three biological replicates for each measurement, and the black horizontal lines indicate the corresponding single predicted value. The error bar range indicates the 95% confidence interval calculated using the standard error of the mean for measurements, and the grey area indicates the 95% confidence interval computed using the infinitesimal jackknife estimate of the standard error for predictions^43^.

Using only the data of six of the seven fluorescence measurements for training – effectively making the model blind for one color – and predicting the DPE of the removed FP results in a mean R^2^ of 0.58 (standard deviation of 0.12) and mean MAE of 0.22 (standard deviation of 0.06) over all seven model predictions for the seven FPs (see **Figure S6**, Supplementary Information). If two of the fluorescence measurements are removed simultaneously (double-color-blinded), the mean R^2^ reduces to 0.53 (standard deviation of 0.13) while the mean MAE remains at 0.22 (standard deviation of 0.04) over all 21 resulting models (see **Figure S7** for up to 6 removed FPs in the training set, Supplementary Information). This suggests that the model is still able to predict DPE of proteins whose sequence data was not part of the training set to an acceptable degree (the MAE reaches 50% of the mean measured DPE).

### Important features determining PE

We investigated the importance of the selected features as indicated by Shapley additive explanation (SHAP) values^44^. The SHAP value is given in the same scale and unit as the model prediction, i.e., DPE in our model, and provides a measure to what extent a feature contributed to the final prediction of the model. One of the advantages of this approach is that SHAP values do not provide only a global measure of feature importance by their mean absolute value, but also a “local” explanation for the features of each individual data point. This means that one can extract information about a feature importance depending on its own magnitude and the magnitude of other features of each individual data point. To determine the SHAP values, all 5 models of the cross-validation were used and their test and training data analyzed.

**Figure 4** shows the 8 most impactful features (highest mean absolute SHAP values) resulting from this analysis. This global feature importance remains consistent among all 5 models whether they predict data from the test or training set (see **Figure 4A**). The inducer concentration appears to have the highest impact. As this feature has a direct influence on the expression level by determining the frequency of initiation of transcription, this is not surprising. The importance is further supported by the SHAP values resembling the typical sigmoidal shape of an induction curve when plotted against increasing inducer concentration (see **Figure 4C**).

**Figure 4.**
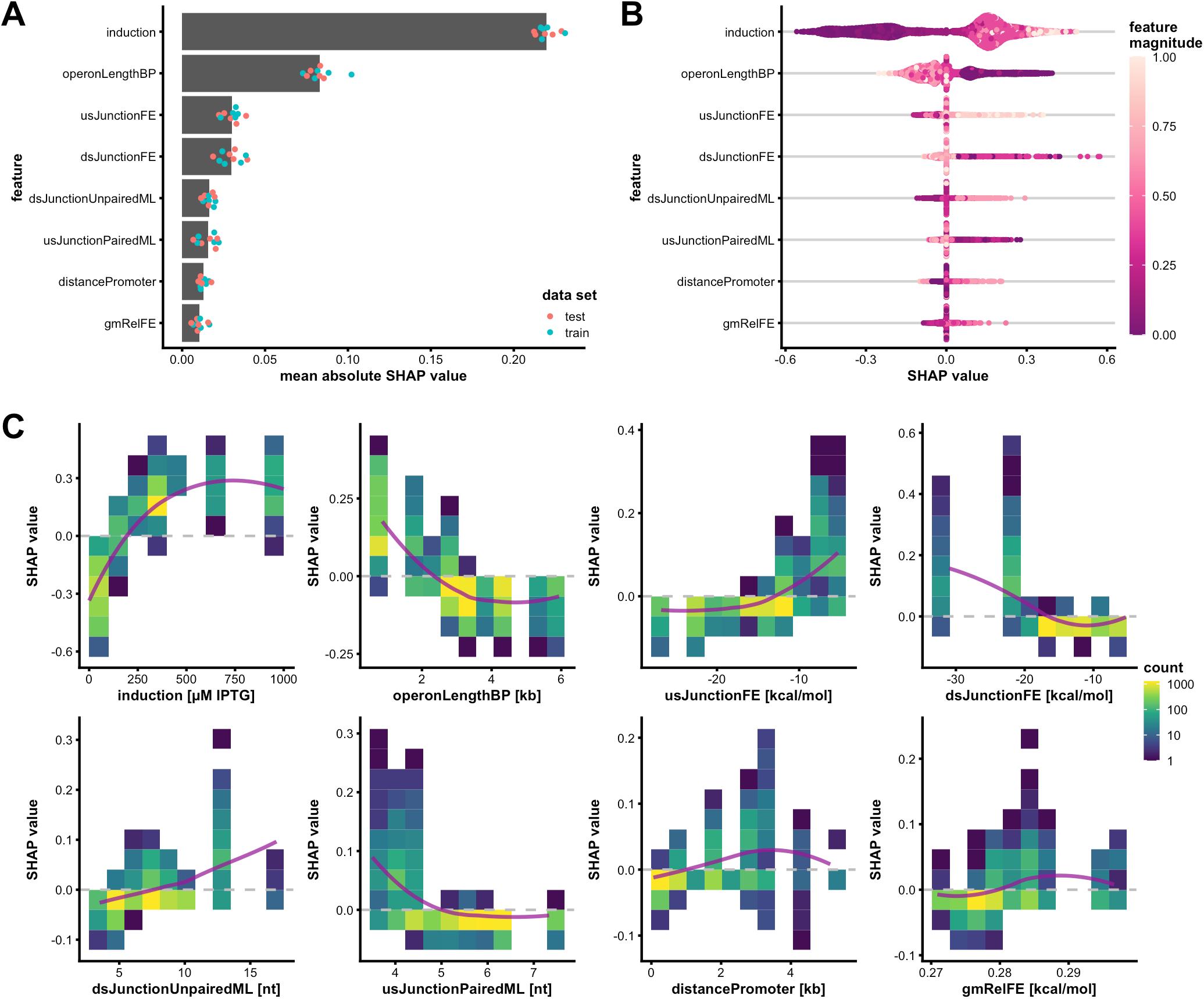
Most impactful features according to SHAP value analysis. For this analysis, all five models of the five-fold cross-validation were used to determine SHAP values using for each model training and test data sets. The features are sorted according to their mean absolute SHAP values and the 8 most impactful (highest overall mean absolute value) features plotted. (**A**) Overall mean absolute SHAP values based on all models of the cross-validation runs (grey bars) and individual mean absolute SHAP values (dots) for the test (red) and training (blue) sets of each model. (**B**) Individual SHAP values for all measurements. Each data point is colored according to each feature’s magnitude (normalized between 0 and 1 for continuous feature values). The data points are plotted as a bee swarm plot to visualize the data density at different SHAP values. (**C**) SHAP values plotted against feature values. The dashed horizontal line marks the position of points with a SHAP value of zero. The colored squares indicate how many values there are in within the indicated area of SHAP values and feature values. The purple line illustrates trends using a LOESS fit over the plotted data. A detailed description corresponding to the feature abbreviations is given in **Table S5** (Supplementary Information).

Two other features and central operon design elements that appeared in our initial qualitative analysis (see **Figure 3A**), namely the operon length (*operonLengthBP)* and the gene’s distance to the promoter (*distancePromoter*), appear among the most impactful features and generally show the same influence on DPE that we observed when averaging our measurements per gene position and operon length. Specifically, increasing operon length leads to decreasing DPE and – within this general trend – increasing distance of a gene from the promoter leads to higher DPE (**Figure 4C**). The general decrease in DPE with operon length can be explained by the increasing likelihood of long sequences to harbor sites that act as terminator during transcription^45,46^ or can be accessed by RNases^47^. On the other hand, the global increase in DPE with increasing distance to the promoter can be seen as an effect of the increased vulnerability of mRNA 5’-end to RNases when transcribed by rapid RNA polymerases like the T7 RNAP^48^. Therefore, increasing distance to the 5’-end increases the chance of shielding structures to appear between the initiation site of degradation and the gene of interest. Against this broader background we observe various patterns in DPE depending on operon context, hence, additional features determine the precise individual DPE for each context. For instance, the junction between a gene of interest and its upstream gene appears to play a particularly important role. In our analysis, we find the following 2 features among the 8 most impactful ones: the RNA secondary structure folding energy of the 100 nucleotides of the upstream junction (*usJunctionFE*) and the mean length of sections with intramolecular paired nucleotides in the RNA structure of the 100 nucleotides of the upstream junction (*usJunctionPairedML*). We clearly see, that with an *usJunctionFE* higher than approx. –15 kcal/mol, representing weaker structures, the SHAP values become strongly positive, suggesting a positive effect on DPE. Similarly, if the *usJunctionPairedML* drops below 5 nucleotides, the SHAP values drastically increase.

We note that this observation is consistent with the influence of two distinct mechanisms described in literature: (i) translational coupling, which contributes to maintaining a specific translation rate as determined by upstream sequences, can be interrupted by increasingly strong structures between genes^21,41^, and (ii) long stretches of paired nucleotides would increase the likelihood of RNase III cleavage at dsRNA, which in turn would reduce the stability of more structured transcript sections^47^.

Also, the downstream junction is represented through 2 features in the list of most impactful features: the RNA secondary structure folding energy of the 100 nucleotides of the downstream junction (*dsJunctionFE*) and the mean length of sections with unpaired nucleotides in the RNA structure of the 100 nucleotides of the downstream junction (*dsJunctionUnpairedML*). In contrast to the upstream junction, here strong structures with less than approx. −20 kcal/mol show highly positive SHAP values. This is consistent with the protective effect that 3’-end structures have shown against 3’ to 5’-directred degradation^40^. Interestingly, for values of *dsJunctionUnpairedML* above 10 nt we find a few strongly positive SHAP values while for values below 5 nt the SHAP values become mostly negative. This is consistent with the observation that long stretches of unpaired nucleotides are associated with an increased vulnerability towards RNases like RNase E^22^. The fact that RNase E effects are probably already captured via the features describing the number of RNase E motifs (*nREmotif, usREmotif*, and *dsREmotif*) might explain why the *dsJunctionUnpairedML* feature does not show any RNase E-associated behavior but instead indirectly describes similar effects as the *usJunctionPairedML* feature and thus long stretches of unpaired nucleotides conversely mean that there are fewer long stretches of paired nucleotides which again would protect against RNase III cleavage.

Lastly, the geometric mean of relative RNA folding energies (absolute values) of all genes in the operon (*gmRelFE*) provides us with information about how strongly the operon is structured on a global scale. While most SHAP values for this feature are relatively low, we observe the most strongly positive SHAP values only for more strongly structured operons (high *gmRelFE*) and the most strongly negative SHAP values only for weaker structures (low *gmRelFE*). This observation is consistent with previous observations that structured RNA can be more resistant against several RNases^40^.

The SHAP value analysis for all remaining features can be seen in **Figure S8** (Supplementary Information). In conclusion, our approach to use a random forest regression model to predict DPE values from sequence features not only results in accurate predictions but also allows extracting feature importance values that align in an interpretable manner consistent with sequence elements individually reported in literature.

## Discussion

In this study, we have presented an unprecedented set of 88 synthetic operons with diverse gene position permutations, including long operons, and simultaneous read out of up to seven reporter proteins per operon. To the best of our knowledge, we present the first study to systematically investigate the effects of operon length and gene order for synthetic operons with more than three genes^38,39^. This is of particular interest for the synthetic biology community as many synthetic pathways and circuits comprise more than three genes^49,50^. Also, natural pathways, for instance in *E. coli*, involve on average 8 steps^51^. Arguably, novel synthetic pathways can be expected to have similar complexity.

Our data, containing 2772 PE measurements, allowed us to investigate the significant effects of gene order and operon length on DPE and to accurately predict the final DPE based on the various sequence features of the individual operon designs using a random forest regression model. Applying this decision tree-based model on our data allowed for a detailed feature importance analysis using SHAP values. This analysis provides valuable insights into which sequence elements have the largest potential of changing expression levels when varied further in future studies and applications.

Using our model, the effect of the composition and arrangement of new synthetic operons on PE levels can be investigated *in silico* before the assembly thus allowing synthetic biologists to make informed and quantitative decisions on operon design candidates at an early stage. For instance, **Figure S10** (Supplementary Information) shows six different arrangements of five FP genes and shows the large context dependency for each of the genes in each operon context. For example, the maximal measured DPE of the same mCardinal gene is 4.45-fold as large as the minimal DPE in different operon arrangements. This difference is also closely captured in the DPE prediction with a factor of 4.4-fold. The other encoded FPs show similar degrees of variation depending on operon context which is largely captured by the model predictions. This example of varying gene arrangement and the resulting large differences in protein expression demonstrates how important it is to not neglect this design decision.

Additionally, the knowledge about the most influential sequence features allows for systematic ‘debugging’ of operon designs that do not fulfill desired specifications in their predicted DPE values by varying these features such as structural elements around gene junctions.

In terms of predicting the DPE of new genes never seen by the model, we consider it an important finding that single and multiple color blinding experiments still resulted in acceptable accuracy. However, the difference in predictive power between the full data set and the “color-blinded” data sets also indicates that the model can be potentially further improved by adding more genes to the measurement set whose feature space further deviates from the currently measured genes (see **Figure S2**, Supplementary Information).

In view of recent successes of machine learning in synthetic biology^52,53^, it is also important to keep a fundamental limitation in mind: the model can only be as good as its training data. The SHAP value analysis reveals that some important features have not been included in our training set with a balanced coverage of values. These features display an uneven distribution over the data set as indicated by gaps in their histograms, for example, *dsJunctionFE* around −30 to −22 kcal/mol and *dsJunctionUnpairedML* around 13 to 17 nt (see **Figure 4C** and **Figure S2C**, Supplementary Information). Therefore, it might be harder to reliably interpret why they appear to be important and in which range of values they are particularly important for the predicted DPE. Additionally, model predictions might be further generalized and the interpretation of these important features could be improved by systematically closing these gaps or extend the range of their values further outside of their limits.

It is also worthwhile to acknowledge that many of the important features might each be a proxy for multiple molecular mechanisms of influence, for example, the intergenic structure can potentially influence transcriptional termination, translational coupling as well as susceptibility to RNases. This provides us with a reminder that biological systems are inherently highly intertwined, complex systems whose accurate prediction requires sufficiently sophisticated models that capture the influence of sequence and systems composition across all each-other-influencing levels from transcriptions over translation to degradation.

Finally, we should point out that our current model is most likely not capable of reliably predicting the PE levels of any arbitrary operon, nor is it intended to. Instead, we focused on a subset of synthetic operons in a highly relevant recombinant system: the T7 expression system^54,55^. As the dynamic formation of the structures that influence translation or RNA degradation is a result of the combined effects of transcription initiation frequency and RNA-synthesis and translation rates, promoter strength, induction, and the choice of the RNA polymerase itself, the importance of certain structural features would potentially change depending on the chosen expression system^56^. We chose the T7 system as it is popular in synthetic biology applications due to the orthogonality of T7 RNAP to endogenous promoters, its high transcription initiation rate, and the high number of developed tools like split T7 RNAP^57^, promoter and RNAP variants with varying activity^58^, and T7 RNAP-dependent plasmid backbones^59^. Since these tools are widely applied and many synthetic operons are constructed using the T7 expression system, our insights will become another valuable tool in the toolbox of T7 systems.

Nevertheless, our approach of combinatorial operon assembly and multiplexed DPE measurement together with machine learning model training is amenable to further extension by, for instance, varying plasmid backbones, promoters or host strains. Thus, synthetic operon predictions could be further generalized to an even larger set of systems applied in synthetic biology.

As we will provide our complete data set and all scripts for model construction and analysis openly upon final publication as well as an early-access version of our user-friendly online tool Ocapy (a tool to predict how <u>O</u>peron <u>C</u>ontext <u>A</u>ffects <u>P</u>rotein expression in s<u>Y</u>nthetic constructs) to predict DPE values (https://bsse.ethz.ch/bpl/software.html) already with this preprint, we hope to provide a foundation for future approaches to design and build new synthetic operons.

## Materials and Methods

### Chemicals and reagents

Unless stated otherwise, chemicals were obtained from Sigma-Aldrich (Buchs, Switzerland). Primers were synthesized by Sigma-Aldrich (see **Table S2** and **S3**, Supplementary Information), and enzymes for standard molecular cloning were obtained from New England Biolabs (Ipswich, USA). If oligos were intended as primers for DNA fragment amplification, they were purchased in 5’-phosphorylated form (see **Table S2**, Supplementary Information) from Integrated DNA Technologies (Coralville, USA). The Ampligase Thermostable DNA Ligase for LCR reaction was obtained from Lucigen (Middleton, USA).

### Strains and cultivation

*Escherichia coli* strain MG1655(DE3) – genotype: K-12 F– *ilvG– rfb-50 rph-1* λ(DE3 [*lacI lacUV5-T7p07 ind1 sam7 nin5]*) (previously constructed^60^) was used for all experiments involving protein expression from introduced plasmids.

The strains were grown in LB Miller broth (Difco™ lysogeny broth (LB)-Miller media, Becton Dickinson (Sparks, USA); LB from hereon) for all initial precultures and cryo-stock preparations. The growth medium for all experiments involving PE and fluorescence measurements was M9 mineral medium^61^ supplemented with TE-US* trace elements^62^ and 0.4% w/v glucose as carbon source (M9G from hereon) and was prepared according to the corresponding literature protocols. Kanamycin (kan) was supplemented at 50 μg mL^-1^.

MG1655(DE3) cryo-stocks containing the plasmids with a particular operon design each were used to first inoculate 200 μL of LB kan in a 96-well F-bottom MTP and incubate it with shaking at 37 °C for 5-7 hours. With 10 μL of this culture, 490 μL of M9G kan in a deep-well block (DWB) were inoculated. The DWB block was sealed with an air-permeable lid (Duetz-system, Kuhner, Switzerland) and incubated overnight at 37 °C and 300 rpm in an incubator appropriately equipped for microtiter plate incubation (Kuhner, Switzerland). The following day, 490 μL of M9G kan in a DWB were inoculated with 10 μL of the M9G overnight culture and incubated for 3 hours at 37 °C and 300 rpm using the Duetz-system incubator. Then the culture was induced using isopropyl β-D-1-thiogalactopyranoside (IPTG) at various concentrations. The induced culture was further incubated for 3 hours at 37 °C and 300 rpm after which samples for measurements were taken (see below).

### Fluorescence measurements via flow cytometry

The fluorescence analysis of the cell suspension was performed using a BD LSR Fortessa SORP (BD Biosciences, Allschwil, Switzerland) equipped with a high-throughput sampling device and 5 lasers (405 nm, 448 nm, 488 nm, 561 nm, and 633 nm). The fluorescence intensities of the different fluorescent proteins were measured using following laser/filter combinations (target FP - excitation wavelength in nm - emission wavelength/bandwidth in nm): mCardinal-633-670/14, mNectarine-561-586/15, LSSmOrange-445-586/15, mNeonGreen-488- 530/11, mT-Sapphire-405-530/30, mTurquoise2-445-473/10, mTagBFP2-405-450/50.

After the measurement, the compensated mean fluorescence value of each channel was extracted using the flow cytometry analysis software FlowJo (version 10.6.2, FlowJo LLC, USA). For the compensation, the monocistronic reference constructs were used to automatically generate a compensation matrix using the FlowJo-built-in compensation function. For each channel, this resulting mean value was divided by the mean fluorescence of the monocistronic reference sample to obtain the DPE value.

### LCR assembly

The LCR bridging oligos were designed as combination of “half-bridges” as described previously in detail^35^. Each half-bridge contains either the sequence of the end or the start of a DNA fragment. While the bridging oligos can contain either the forward or the reverse strand sequence, in this study they were designed to contain only the forward sequence according to the direction of the encoded operon genes. Each half bridge was chosen to be just long enough – starting from the junction – to have a melting temperature of approximately 70 °C as estimated by the OligoAnalyzer online tool (version 3.1, https://eu.idtdna.com/calc/analyzer) with an oligo concentration of 0.03 μM, a sodium ion concentration of 50 mM, and a magnesium ion concentration of 10 mM. The resulting fragment-end half-bridges and fragment-start half-bridges were then joined to form full bridging oligos that guide the ligation of the desired fragment combination (see **Table S3** for bridging oligo sequences, Supplementary Information).

To perform the assembly, the required DNA fragments were prepared with a concentration of 75 nM and appropriate bridging oligos were diluted to 750 nM. Per LCR, 1 μL of each required DNA fragment and corresponding bridging oligo were added to 2.5 μL of Ampligase 10X Reaction Buffer (Lucigen, USA), 2.25 μL of 5 M betaine, 2 μL of DMSO, and 1.5 μL of 5 U μL^-1^ Ampligase Thermostable DNA Ligase (Lucigen, USA). Water was added to a total volume of 25 μL. This results in a final DNA fragment concentration of 3 nM and bridging oligo concentration of 30 nM each in an LCR. The thermocycler protocol was as follows: 120 s at 94 °C initial denaturation, 50 cycles of 10 s at 94 °C, 30 s at 55 °C, and 60 s at 66 °C, and a final hold step at 4 °C. Three of the assembled fragments in each reaction were backbone fragments (fDG163, fDG164, and fDG197; see **Table S1**, Supplementary Information) which resulted in full plasmids after assembly. The LCR product was directly added to chemically competent MG1655(DE3) cells for transformation. The transformation mix was plated on LB agar supplemented with the appropriate antibiotic.

### Data set processing

The 2772 fluorescence measurements (DPE values) of individual FPs were matched with information on the unique operon for which they were measured, the relevant inducer concentration, and the experimental replicate, and then assigned to the 19 features listed in **Table S5**. Sequence-dependent features for each operon were extracted from corresponding operon sequences using the *R*^63^ packages *Biostrings*^64^ and *XNAString*^65^ and assigned to each corresponding measurement.

### Random forest model

All random forest models were built in *R* using the package *ranger*^66^. The 19 features listed in **Table S5** (Supplementary Information) were used as input variables and the measured DPE as dependent variable for the regression. For the 5-fold cross-validation, the data set was stratified to ensure that biological replicates could only be either in the training set or in the test set, resulting in 789 groups of biological replicates. These groups were randomly split into 5 subsets for cross-validation. Within each cross-validation run, variables containing empty values, since they are undefined at the edges of the operon, i.e. undefined upstream or downstream values for *usRBS, usREMotif*, and *dsREMotif*, were imputed using the package *missRanger*,^67^ for each test and training set separately. Hyperparameters were learned for each training set separately using the package *tuneRanger*,^68^ with standard settings.

For the “color-blinded” experiments, all data rows containing measurements of one (for 1-blinded) or several (for 2- to 6-blinded) FPs were removed from the training set and defined as test set. All subsequent steps to train and test the model were equivalent to the 5-fold crossvalidation.

Model performance was measured using the coefficient of determination (R^2^) and the mean absolute error (MAE) between the measured and predicted DPE.

### SHAP value analysis

The importance of each feature for the random forest model prediction was determined using the Shapley additive explanation (SHAP) values^44^ and the corresponding *R* package implementation *shapper*^69^. The SHAP values were calculated for each of the five training and test set resulting from cross-validation.

## End Matter

### Author Contributions and Notes

D. G. and S. P. conceived the study and planned experiments. D. G. performed experiments and computational works. D. G., N. B., and S. P. analyzed the data. S. P. coordinated the study. D. G. and S. P. wrote the manuscript, and all authors reviewed and edited it.

The authors declare no competing interests.

This article contains supporting information online.

### Data availability

Individual DPE measurements, corresponding feature values, and scripts for data processing, model training, and plotting will be made available upon final publication.

## Acknowledgements

The authors would like to acknowledge expert support from Gregor Schmidt from the D-BSSE Lab Automation Facility and Verena Jäggin from the Sigle Cell Facility, and very helpful discussions on random forest models with Susan Posada and on multiple protein systems with Johannes Haerle.

## Funding

This work was supported by the European Commission [project ST-FLOW, grant number 289326] and the Swiss National Science Foundation under the NCCR “Molecular Systems Engineering”.

## Notes

### Competing Interest Statement

The authors have declared no competing interest.

## References

1. Clarke, L. & Kitney, R. Developing synthetic biology for industrial biotechnology applications. Biochem Soc T 48, 113–122 (2020).

2. Karoui, M. E., Hoyos-Flight, M. & Fletcher, L. Future Trends in Synthetic Biology—A Report. Frontiers Bioeng Biotechnology 7, 175 (2019).

3. Hold, C., Billerbeck, S. & Panke, S. Forward design of a complex enzyme cascade reaction. Nat Commun 7, 12971 (2016).

4. Ro, D.-K. et al. Production of the antimalarial drug precursor artemisinic acid in engineered yeast. Nature 440, 940–943 (2006).

5. Elowitz, M. B. & Leibler, S. A synthetic oscillatory network of transcriptional regulators. Nature 403, 335–338 (2000).

6. Ausländer, S., Ausländer, D., Müller, M., Wieland, M. & Fussenegger, M. Programmable single-cell mammalian biocomputers. Nature 487, 123–127 (2012).

7. Jeschek, M., Gerngross, D. & Panke, S. Rationally reduced libraries for combinatorial pathway optimization minimizing experimental effort. Nat Commun 7, 11163 (2016).

8. Ng, C. Y., Farasat, I., Maranas, C. D. & Salis, H. M. Rational design of a synthetic Entner–Doudoroff pathway for improved and controllable NADPH regeneration. Metab Eng 29, 86–96 (2015).

9. Hwang, H. J., Lee, S. Y. & Lee, P. C. Engineering and application of synthetic nar promoter for fine-tuning the expression of metabolic pathway genes in Escherichia coli. Biotechnol Biofuels 11, 103 (2018).

10. Exley, K. et al. Utilising datasheets for the informed automated design and build of a synthetic metabolic pathway. J Biol Eng 13, 8 (2019).

11. Hossain, A. et al. Automated design of thousands of nonrepetitive parts for engineering stable genetic systems. Nat Biotechnol 38, 1466–1475 (2020).

12. Shis, D. L., Hussain, F., Meinhardt, S., Swint-Kruse, L. & Bennett, M. R. Modular, Multi-Input Transcriptional Logic Gating with Orthogonal LacI/GalR Family Chimeras. Acs Synth Biol 3, 645–651 (2014).

13. Jacob, F., Perrin, D., Sanchez, C. & Monod, J. [Operon: a group of genes with the expression coordinated by an operator]. Comptes rendus hebdomadaires des seances de l’Academie des sciences 250, 1727–1729 (1960).

14. Salgado, H., Moreno-Hagelsieb, G., Smith, T. F. & Collado-Vides, J. Operons in Escherichia coli: Genomic analyses and predictions. Proc National Acad Sci 97, 6652–6657 (2000).

15. Rennig, M. et al. Industrializing a Bacterial Strain for l-Serine Production through Translation Initiation Optimization. Acs Synth Biol 8, 2347–2358 (2019).

16. Nowroozi, F. F. et al. Metabolic pathway optimization using ribosome binding site variants and combinatorial gene assembly. Appl Microbiol Biot 98, 1567–1581 (2014).

17. Kelwick, R. et al. A Forward-Design Approach to Increase the Production of Poly-3-Hydroxybutyrate in Genetically Engineered Escherichia coli. Plos One 10, e0117202 (2015).

18. Siqueira, G. M. V. de, Silva-Rocha, R. & Guazzaroni, M.-E. Turning the Screw: Engineering Extreme pH Resistance in Escherichia coli through Combinatorial Synthetic Operons. Acs Synth Biol 9, 1254–1262 (2020).

19. Nishizaki, T., Tsuge, K., Itaya, M., Doi, N. & Yanagawa, H. Metabolic engineering of carotenoid biosynthesis in Escherichia coli by ordered gene assembly in Bacillus subtilis. Applied and Environmental Microbiology 73, 1355–1361 (2007).

20. Rex, G., Surin, B., Besse, G., Schneppe, B. & McCarthy, J. E. The mechanism of translational coupling in Escherichia coli. Higher order structure in the atpHA mRNA acts as a conformational switch regulating the access of de novo initiating ribosomes. 269, 18118–18127 (1994).

21. Huber, M. et al. Translational coupling via termination-reinitiation in archaea and bacteria. Nat Commun 10, 4006 (2019).

22. Mackie, G. A. RNase E: at the interface of bacterial RNA processing and decay. Nat Rev Microbiol 11, 45–57 (2013).

23. Conway, T. et al. Unprecedented High-Resolution View of Bacterial Operon Architecture Revealed by RNA Sequencing. Mbio 5, e01442–14 (2014).

24. Mao, X. et al. Revisiting operons: an analysis of the landscape of transcriptional units in E. coli. Bmc Bioinformatics 16, 356 (2015).

25. Chu, J. et al. Non-invasive intravital imaging of cellular differentiation with a bright red-excitable fluorescent protein. Nat Methods 11, 572–578 (2014).

26. Johnson, D. E. et al. Red Fluorescent Protein pH Biosensor to Detect Concentrative Nucleoside Transport. J Biol Chem 284, 20499–20511 (2009).

27. Shcherbakova, D. M., Hink, M. A., Joosen, L., Gadella, T. W. J. & Verkhusha, V. V. An Orange Fluorescent Protein with a Large Stokes Shift for Single-Excitation Multicolor FCCS and FRET Imaging. J Am Chem Soc 134, 7913–7923 (2012).

28. Shaner, N. C. et al. A bright monomeric green fluorescent protein derived from Branchiostoma lanceolatum. Nat Methods 10, 407–409 (2013).

29. Ai, H.-W., Hazelwood, K. L., Davidson, M. W. & Campbell, R. E. Fluorescent protein FRET pairs for ratiometric imaging of dual biosensors. Nature methods 5, 401–403 (2008).

30. Goedhart, J. et al. Structure-guided evolution of cyan fluorescent proteins towards a quantum yield of 93%. Nat Commun 3, 751 (2012).

31. Subach, O. M., Cranfill, P. J., Davidson, M. W. & Verkhusha, V. V. An enhanced monomeric blue fluorescent protein with the high chemical stability of the chromophore. PLoS One 6, e28674 (2011).

32. Saunders, J. R., Pickup, R. W., Morgan, J. A., Winstanley, C. & Saunders, V. A. Molecular Microbial Ecology Manual. in 193–204 (1996). doi:10.1007/978-94-009-0215-2_12.

33. Stahl, S. W. et al. Single-molecule dissection of the high-affinity cohesin-dockerin complex. Proceedings of the National Academy of Sciences 109, 20431–20436 (2012).

34. Song, C. W., Kim, J. W., Cho, I. J. & Lee, S. Y. Metabolic Engineering of Escherichia coli for the Production of 3-Hydroxypropionic Acid and Malonic Acid through β-Alanine Route. Acs Synth Biol 5, 1256–1263 (2016).

35. Kok, S. de et al. Rapid and Reliable DNA Assembly via Ligase Cycling Reaction. Acs Synth Biol 3, 97–106 (2014).

36. Salis, H. M., Mirsky, E. A. & Voigt, C. A. Automated Design of Synthetic Ribosome Binding Sites to Precisely Control Protein Expression. Nat Biotechnol 27, 946–950 (2009).

37. Martínez-García, E. et al. SEVA 3.0: an update of the Standard European Vector Architecture for enabling portability of genetic constructs among diverse bacterial hosts. Nucleic Acids Res 48, 3395–3395 (2020).

38. Lim, H. N., Lee, Y. & Hussein, R. Fundamental relationship between operon organization and gene expression. Proc National Acad Sci 108, 10626–10631 (2011).

39. Chizzolini, F., Forlin, M., Cecchi, D. & Mansy, S. S. Gene Position More Strongly Influences Cell-Free Protein Expression from Operons than T7 Transcriptional Promoter Strength. Acs Synth Biol 3, 363–371 (2014).

40. Dar, D. & Sorek, R. Extensive reshaping of bacterial operons by programmed mRNA decay. Plos Genet 14, e1007354 (2018).

41. Osterman, I. A., Evfratov, S. A., Sergiev, P. V. & Dontsova, O. A. Comparison of mRNA features affecting translation initiation and reinitiation. Nucleic Acids Res 41, 474–486 (2013).

42. Cetnar, D. P. & Salis, H. M. Systematic Quantification of Sequence and Structural Determinants Controlling mRNA stability in Bacterial Operons. Acs Synth Biol 10, 318–332 (2021).

43. Wager, S., Hastie, T. & Efron, B. Confidence Intervals for Random Forests: The Jackknife and the Infinitesimal Jackknife. J Mach Learn Res Jmlr 15, 1625–1651 (2014).

44. Lundberg, S. M. et al. From local explanations to global understanding with explainable AI for trees. Nat Mach Intell 2, 56–67 (2020).

45. Ray-Soni, A., Bellecourt, M. J. & Landick, R. Mechanisms of Bacterial Transcription Termination: All Good Things Must End. Annu Rev Biochem 85, 1–29 (2016).

46. Chen, Y.-J. et al. Characterization of 582 natural and synthetic terminators and quantification of their design constraints. Nat Methods 10, 659–664 (2013).

47. Hui, M. P., Foley, P. L. & Belasco, J. G. Messenger RNA Degradation in Bacterial Cells. Annu Rev Genet 48, 1–23 (2014).

48. Makarova, O. V., Makarov, E. M., Sousa, R. & Dreyfus, M. Transcribing of Escherichia coli genes with mutant T7 RNA polymerases: Stability of lacZ mRNA inversely correlates with polymerase speed. Proceedings of the National Academy of Sciences 92, 12250–12254 (1995).

49. Smanski, M. J. et al. Functional optimization of gene clusters by combinatorial design and assembly. Nat Biotechnol 32, 1241–1249 (2014).

50. Bahls, M. O., Platz, L., Morgado, G., Schmidt, G. W. & Panke, S. Directed evolution of biofuel-responsive biosensors for automated optimization of branched-chain alcohol biosynthesis. Metab Eng 69, 98–111 (2022).

51. Arita, M. The metabolic world of Escherichia coli is not small. P Natl Acad Sci Usa 101, 1543–1547 (2004).

52. Volk, M. J. et al. Biosystems Design by Machine Learning. Acs Synth Biol 9, 1514–1533 (2020).

53. Lawson, C. E. et al. Machine learning for metabolic engineering: A review. Metab Eng 63, 34–60 (2021).

54. Wang, W. et al. Bacteriophage T7 transcription system: an enabling tool in synthetic biology. Biotechnol Adv 36, 2129–2137 (2018).

55. Shis, D. L. & Bennett, M. R. Synthetic biology: the many facets of T7 RNA polymerase. 745–745 (2014).

56. Pan, T. & Sosnick, T. RNA FOLDING DURING TRANSCRIPTION. vol. 35 (2006).

57. Shis, D. L. & Bennett, M. R. Library of synthetic transcriptional AND gates built with split T7 RNA polymerase mutants. Proc National Acad Sci 110, 5028–5033 (2013).

58. Temme, K., Hill, R., Segall-Shapiro, T. H., Moser, F. & Voigt, C. A. Modular control of multiple pathways using engineered orthogonal T7 polymerases. Nucleic Acids Res 40, 8773–8781 (2012).

59. Becker, K., Meyer, A., Roberts, T. M. & Panke, S. Plasmid replication based on the T7 origin of replication requires a T7 RNAP variant and inactivation of ribonuclease H. Nucleic Acids Res 49, 8189–8198 (2021).

60. Napiorkowska, M., Pestalozzi, L., Panke, S., Held, M. & Schmitt, S. High-Throughput Optimization of Recombinant Protein Production in Microfluidic Gel Beads. Small 17, 2005523 (2021).

61. Sambrook, J. F. & Russell, D. W. Molecular Cloning: A Laboratory Manual. (Cold Spring Harbor Laboratory Press, 2001).

62. Panke, S., Lorenzo, V. de, Kaiser, A., Witholt, B. & Wubbolts, M. G. Engineering of a Stable Whole-Cell Biocatalyst Capable of (S)-Styrene Oxide Formation for Continuous Two-Liquid-Phase Applications. Appl Environ Microb 65, 5619–5623 (1999).

63. R Core Team. R: A Language and Environment for Statistical Computing. (R Foundation for Statistical Computing, 2020).

64. Pagès, H., Aboyoun, P., Gentleman, R. & DebRoy, S. Biostrings: Efficient manipulation of biological strings. R package version 2.60.2. (2021).

65. Górska, A. et al. XNAString: Efficient Manipulation of Modified Oligonucleotide Sequences. R package version 1.0.2. (2021).

66. Wright, M. N. & Ziegler, A. ranger : A Fast Implementation of Random Forests for High Dimensional Data in C++ and R. J Stat Softw 77, (2017).

67. Mayer, M. missRanger: Fast Imputation of Missing Values. R package version 2.1.3. (2021).

68. Probst, P., Wright, M. N. & Boulesteix, A. Hyperparameters and tuning strategies for random forest. Wiley Interdiscip Rev Data Min Knowl Discov 9, 281 (2019).

69. Maksymiuk, S., Gosiewska, A. & Biecek, P. shapper: Wrapper of Python Library “shap”. R package version 0.1.3. (2020)

