## Supplementary Information for "Systematic investigation of synthetic operon designs enables prediction and control of expression levels of multiple proteins"

**This PDF file includes:**

Figure S1 to S10
Table S1 to S5

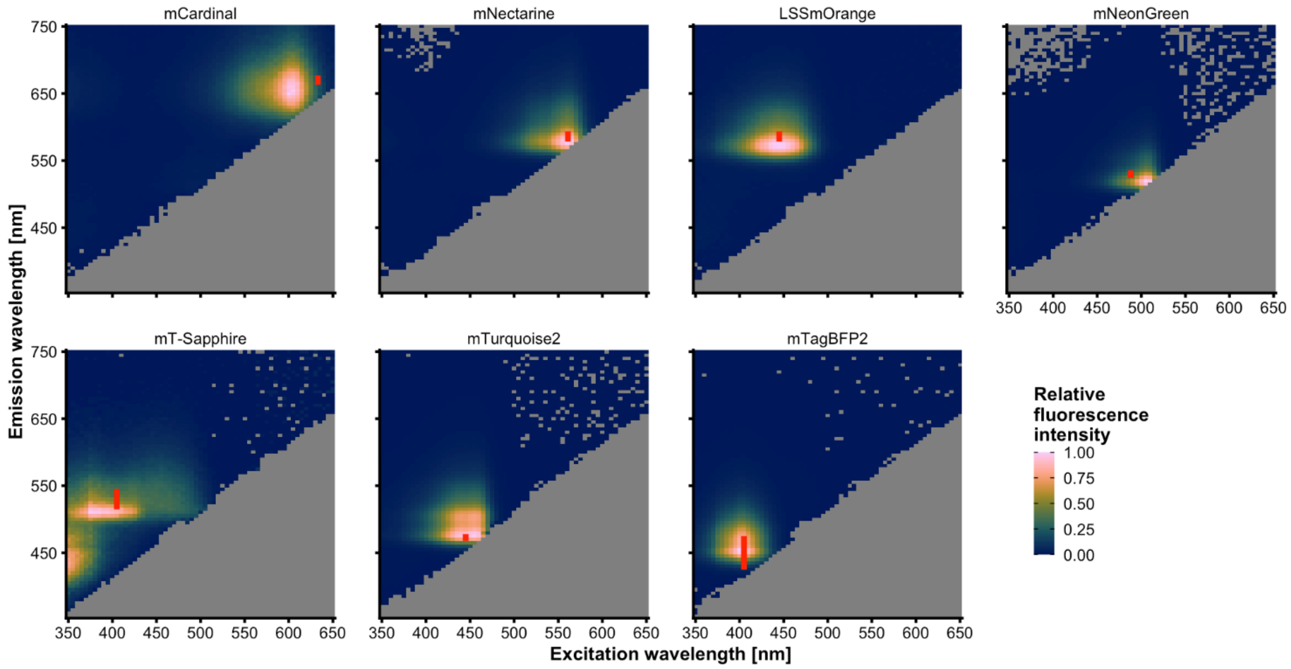


**Figure S1. Fluorescence spectra of reporter proteins.** The seven fluorescent proteins (FPs) mCardinal, mNectarine, LSSmOrange, mNeonGreen, mT-Sapphire, mTurquoise2, and mTagBFP2 were expressed and measured in individual *E. coli* cultures. The color indicates the measured fluorescence intensity relative to maximum fluorescence of each FP. Grey areas indicate emission/excitation combinations that were not measured (lower-right triangle) or measurements that resulted in negative values after background correction (scattered grey spots in the upper area). The red line segment in each 2D spectrum indicates emission and excitation parameters used in flow cytometry measurements of each FP. The line's position on the excitation wavelength axis indicates the wavelength of the laser used for excitation. The range of the line in the vertical direction indicates the wavelength range of the emission band-pass filter used for the measurement of the corresponding FP.

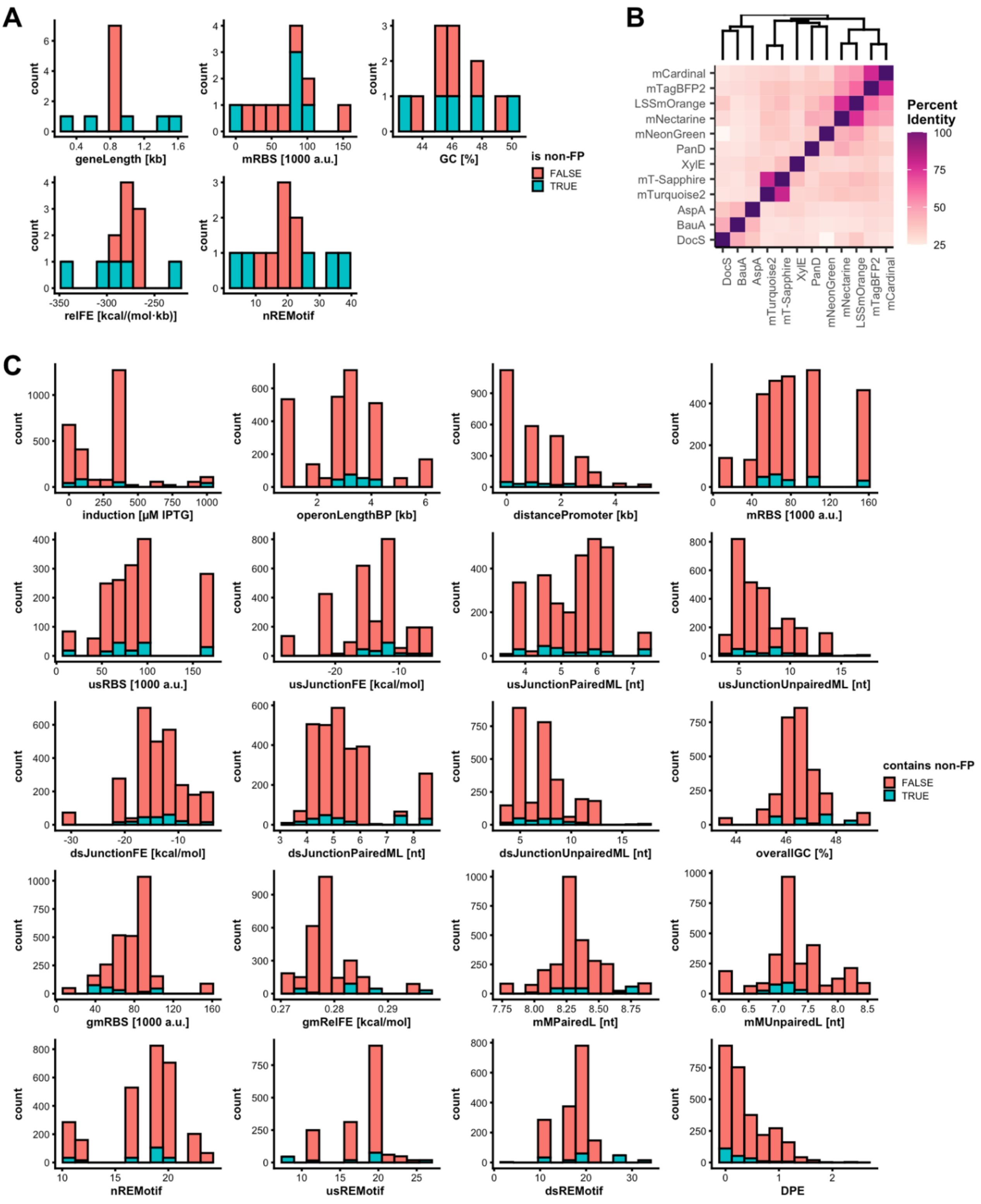


**Figure S2. Diversity of investigated genes and operons.** (**A**) Distribution of the gene-specific features that were used directly (*mRBS*, *nREMotif*) or indirectly (*geneLength*→*operonLengthBP*, GC→*overallGC*, *relFE*→*gmRelFE*) for training the random forest models (see **Table S5**, Supplementary Information). Parts of the distributions that are contributed by the genes coding for fluorescent proteins (FP) or non-fluorescent proteins (non-FP) are marked in red and blue respectively. (**B**) Identity matrix of the genes coding for the listed proteins. The percent identity is based on a Clustal Omega sequence alignment. (**C**) Distribution of all features used for training the random forest models in the complete data set. Additionally, the distribution of all measured DPE values is plotted in the last panel. Parts of the distributions that are contributed by operons containing only genes coding for FPs or operons also containing genes coding for non-FPs are marked in red and blue respectively.

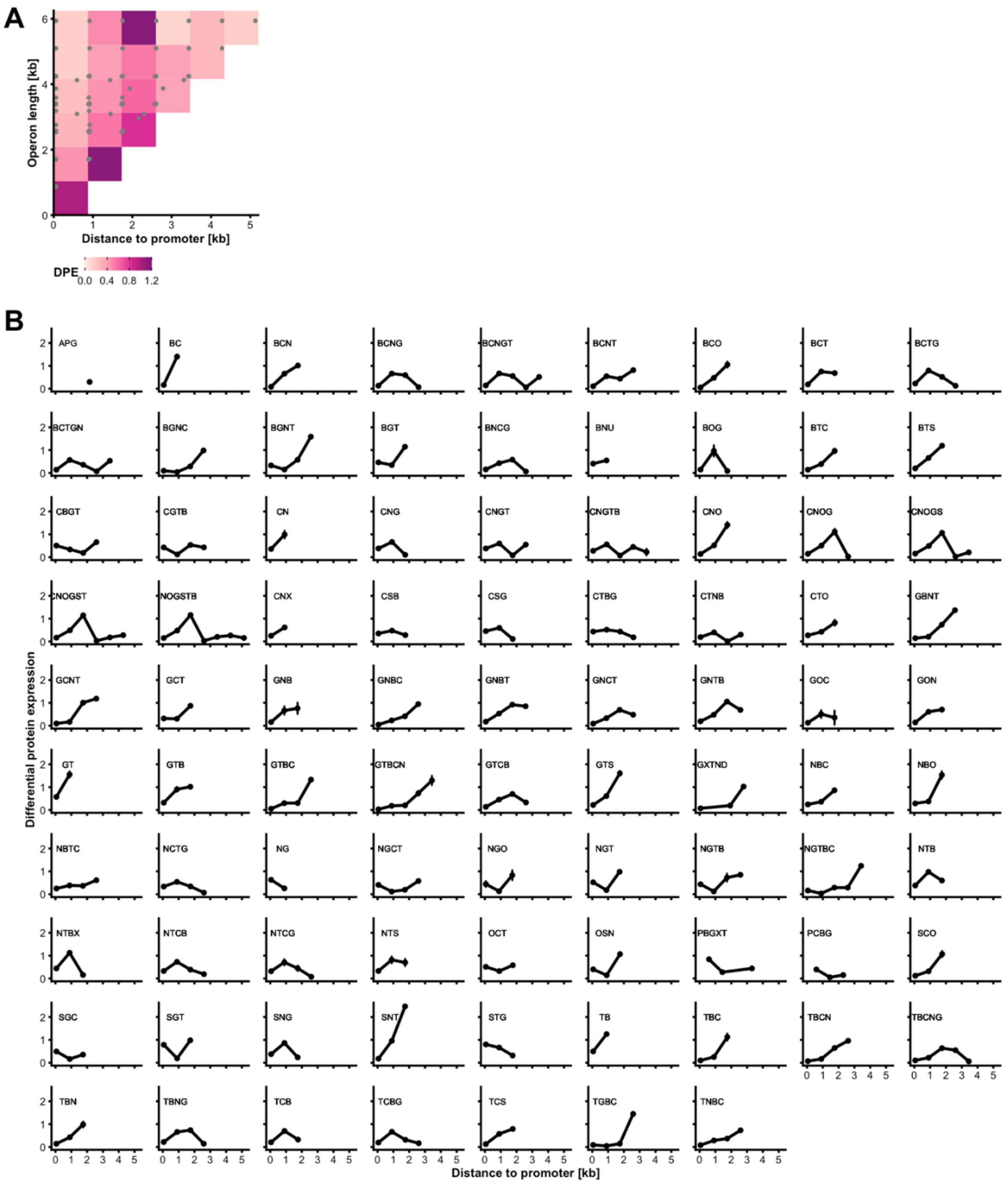


**Figure S3. DPE as a function of operon length, gene distance to promoter, and gene order for all 88 constructed operon designs.** (**A**) Summary DPE matrix for all operons constructs (88 operons designs) and monocistronic references. The horizontal axis indicates the distance of a gene's start codon to the transcription start site (TSS) in kb. The vertical axis indicates the operon length between TSS and terminator in kb. The color of each tile indicates the mean DPE value of all encoded proteins whose genes fall within its boundaries. The bottom left tile contains only the monocistronic reference constructs and has therefore per definition a value of 1. In contrast to **Figure 2A** this plot contains two additional operons with an operon length of more than 4.5 kb. Therefore, the most upper row of tiles contains the only heptacistronic operon design CNOGSTB. The underlying individual operon length and promoter distance values of each operon design are indicated by grey points. (**B**) PE patterns of all 88 constructed bi- to heptacistronic operon designs. Each panel represents one operon design indicated by the letters above each graph (C: mCardinal, N: mNectarine, O: LSSmOrange, G: mNeonGreen, S: mT-Sapphire, T: mTurquoise2, B: mTagBFP2, X: XylE, D: DocS, A: AspA, P: PanD, U: BauA). For instance, the operon design CSG codes for three FPs in following order: mCardinal, mT-Sapphire, mNeonGreen. Only the points for FPs are plotted. The vertical axis indicates the DPE. The horizontal axis indicates the respective gene's start codon distance to the TSS in kb. The line connecting the points illustrates the expression level pattern across each operon (non-FPs are skipped). The points indicate the mean value of at least three biological replicates for each measurement. The error bars indicate one standard deviation of the mean in each direction. The inducer concentration in all plotted experiments was 400 µM IPTG.

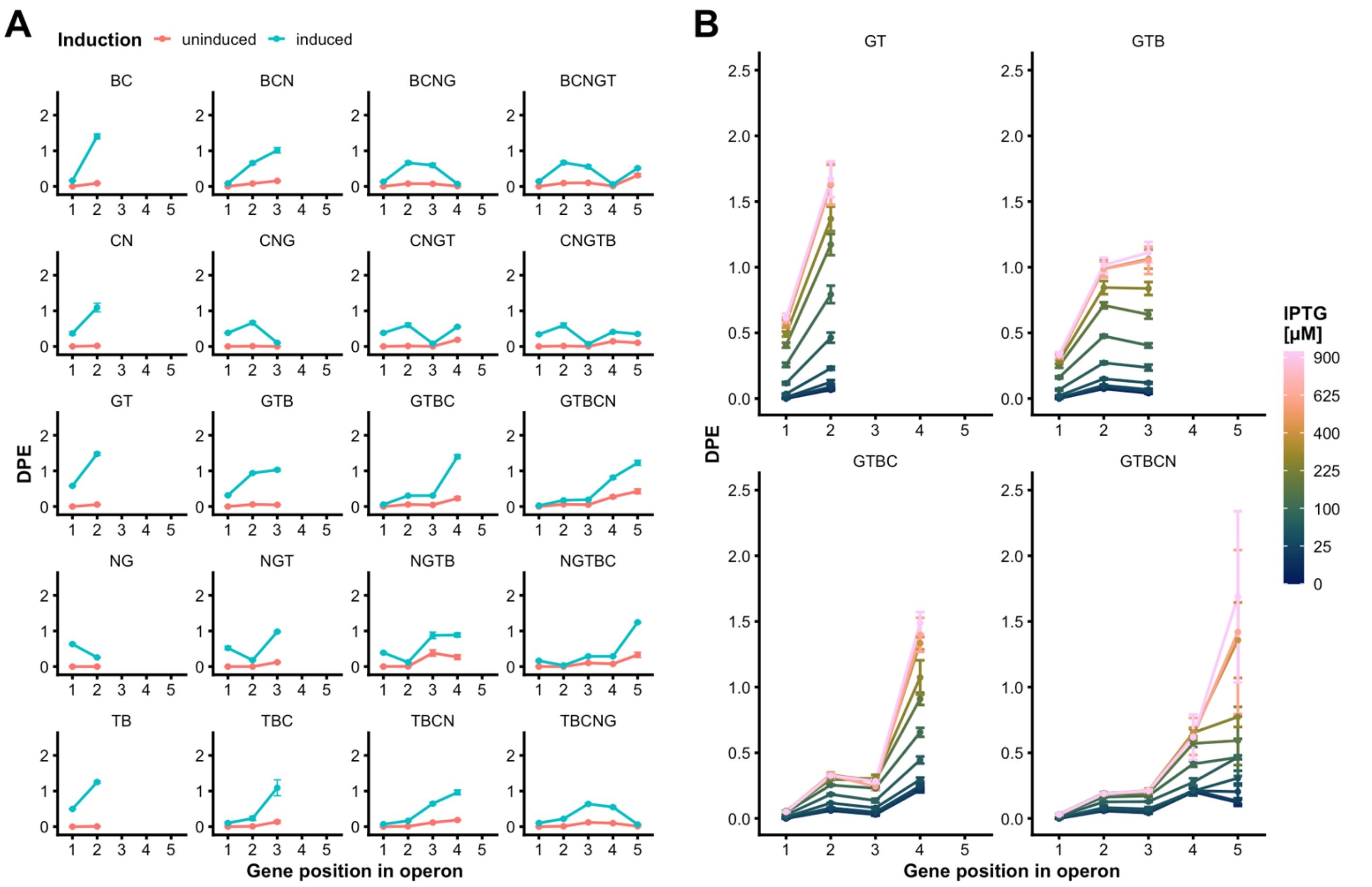


**Figure S4. Effect of induction on DPE for genes with different promoter distances.** (**A**) Expression level of 20 bi- to pentacistronic operon designs with (blue) and without (red) inducer. Each operon is plotted in a separate panel. The letters above each panel represent the gene order in each operon. The vertical axes indicate the DPE. The horizontal axes indicate the position of a gene within an operon. The inducer concentration for induced samples in this experiment was 0 or 400 µM IPTG. (**B**) Expression level of a series bi- to pentacistronic operons (GT, GTB, GTBC, and GTBCN) with varying IPTG concentrations. The color gradient indicates different IPTG concentrations and is scaled using the square root of the concentration for improved visualization. The points indicate the mean value of three biological replicates for each measurement. The error bars indicate one standard deviation of the mean in each direction.

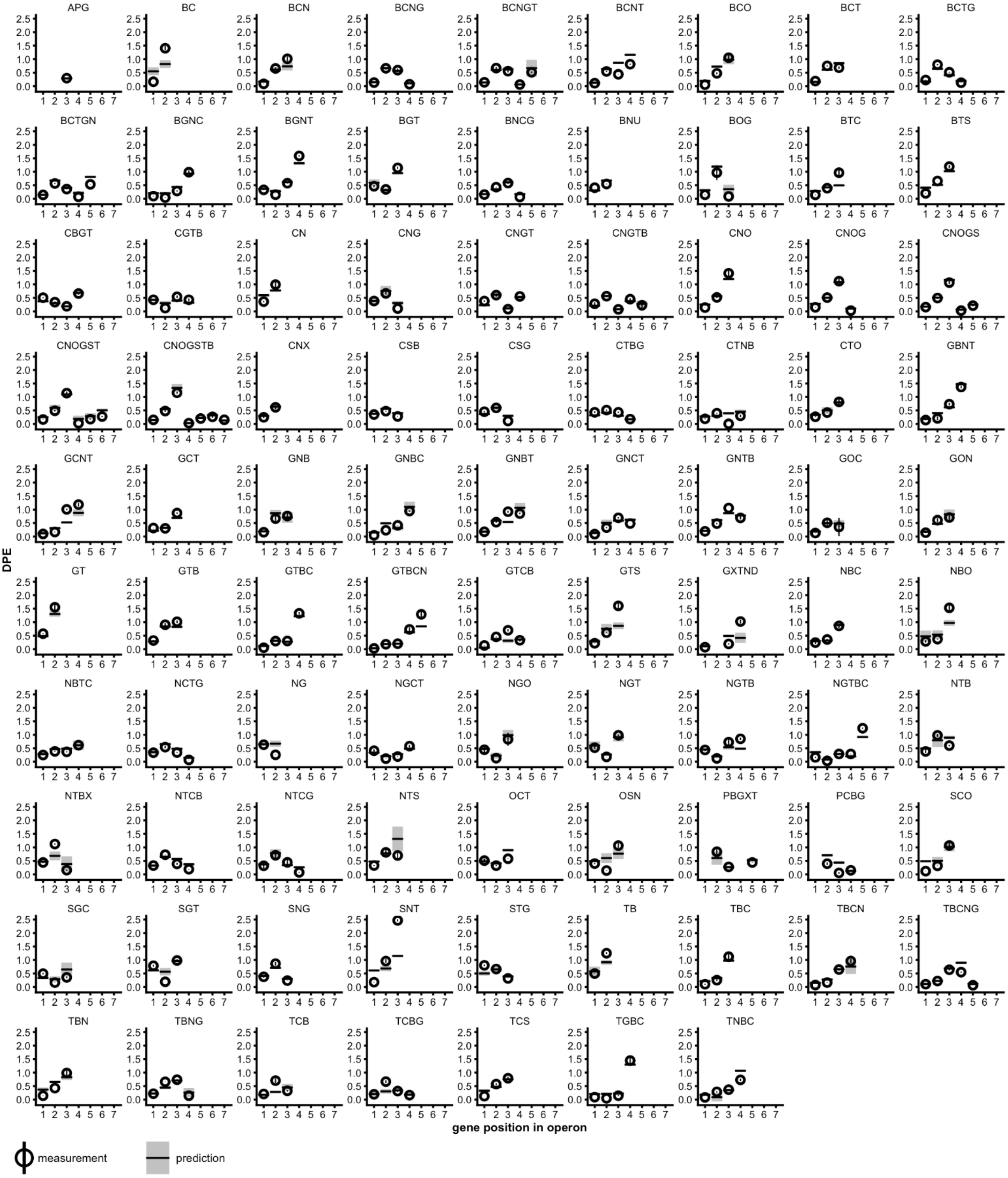


**Figure S5. Random forest regression model-based DPE prediction compared measurements.** The measured DPE and the predicted DPE for all 88 operon designs at an inducer concentration of 400 μM IPTG used in the five test sets of a 5-fold cross-validation of the model (see **Figure 3**) are plotted as circle with error bar and as horizontal line with grey area respectively. Each panel represents one operon design indicated by the order of the letters above each panel (C: mCardinal, N: mNectarine, O: LSSmOrange, G: mNeonGreen, S: mT-Sapphire, T: mTurquoise2, B: mTagBFP2, X:XylE, D:DocS, A:AspA, P:PanD, U:BauA). The circles indicate the mean value of at least three biological replicates for each measurement, and the black horizontal lines indicate the corresponding single predicted value. The error bar range indicates the 95% confidence interval calculated using the standard error of the mean for measurements, and the grey area indicates the 95% confidence interval computed using the infinitesimal jackknife estimate of the standard error for predictions[43]. For X, D, A, P, and U no values are plotted as these non-FPs could not be measured in the experimental setup.

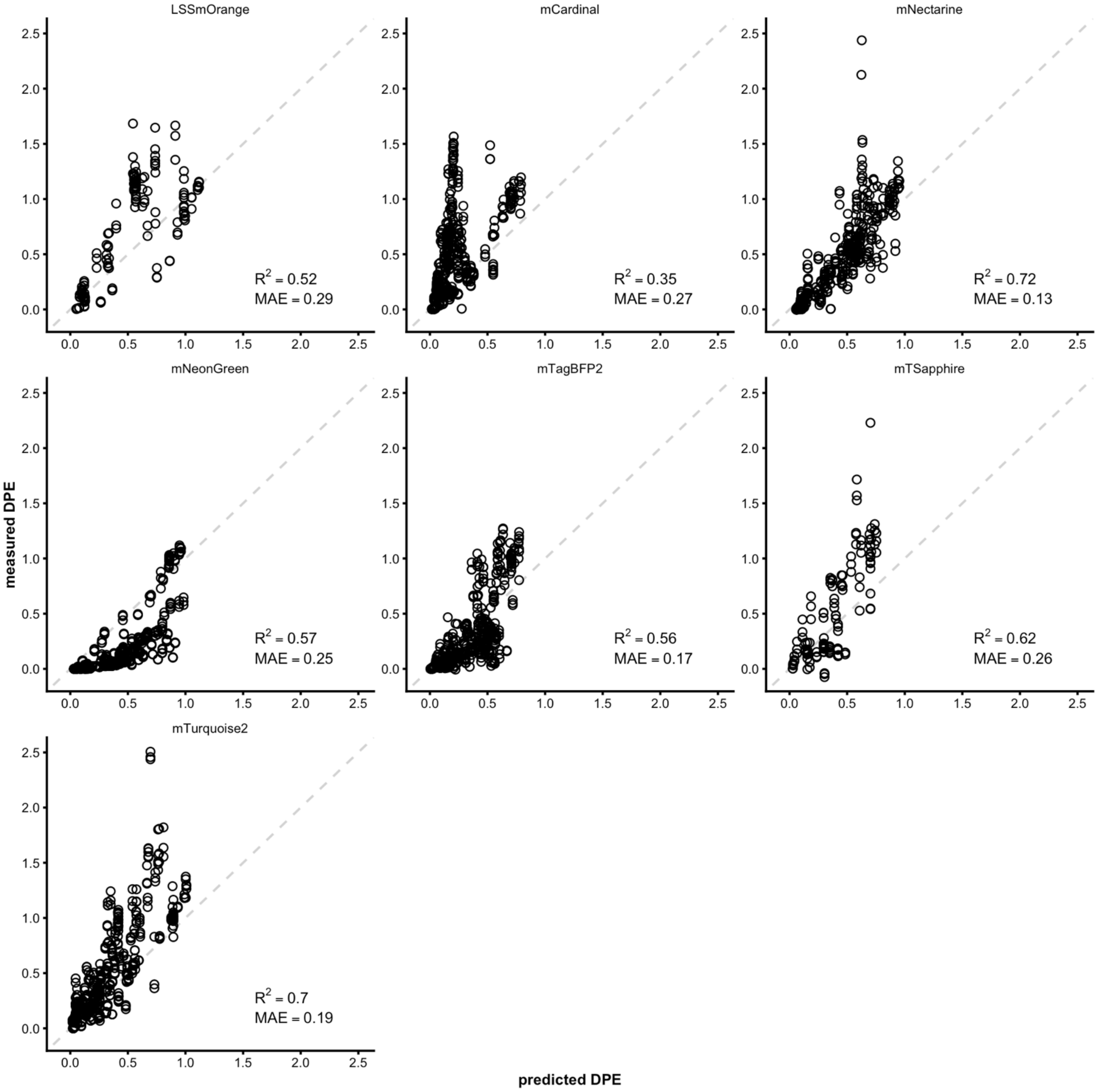


**Figure S6. “Color-blinded” model cross-validation.** Measured DPE plotted against the predicted DPE of seven different random forest regression models. The random forest models were trained on all operon designs measured in this study. However, for each model, the data rows of one of the measured FPs were removed before training. The test set for each model consisted of the data for the corresponding removed FP data. Only the predictions and measurements for each test set are plotted. The FP name above the panels is the FP of which the data was hidden for model training and then tested to generate the here depicted plots. Each panel also contains the coefficient of determination ($R^2$) and mean absolute error (MAE) for each model test. The dashed diagonal line indicates the optimal position of points where a prediction would exactly match the measurement.

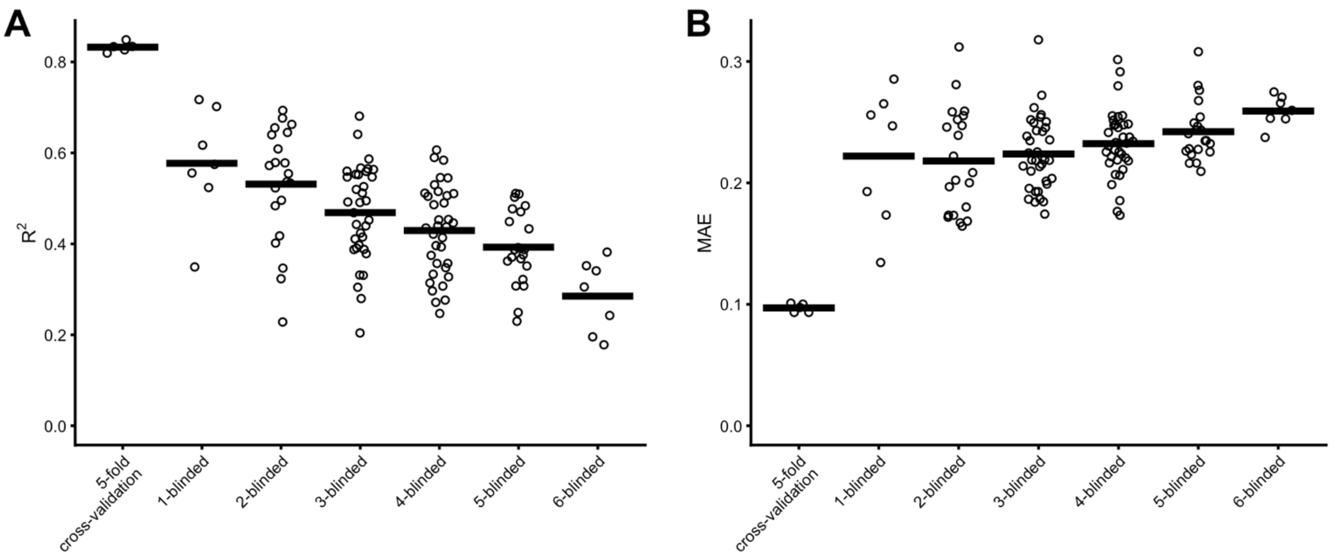


**Figure S7. Cross-validation comparison.** Coefficient of determination ($R^2$; **A**) and mean absolute error (MAE, **B**) of the 5-fold cross-validation (see **Figure 3**) and fluorescence data row "blinding" cross-validation with increasing number of simultaneously removed fluorescence data from training set (see also **Figure S6**). The horizontal bars indicate the mean values within a cross-validation set.

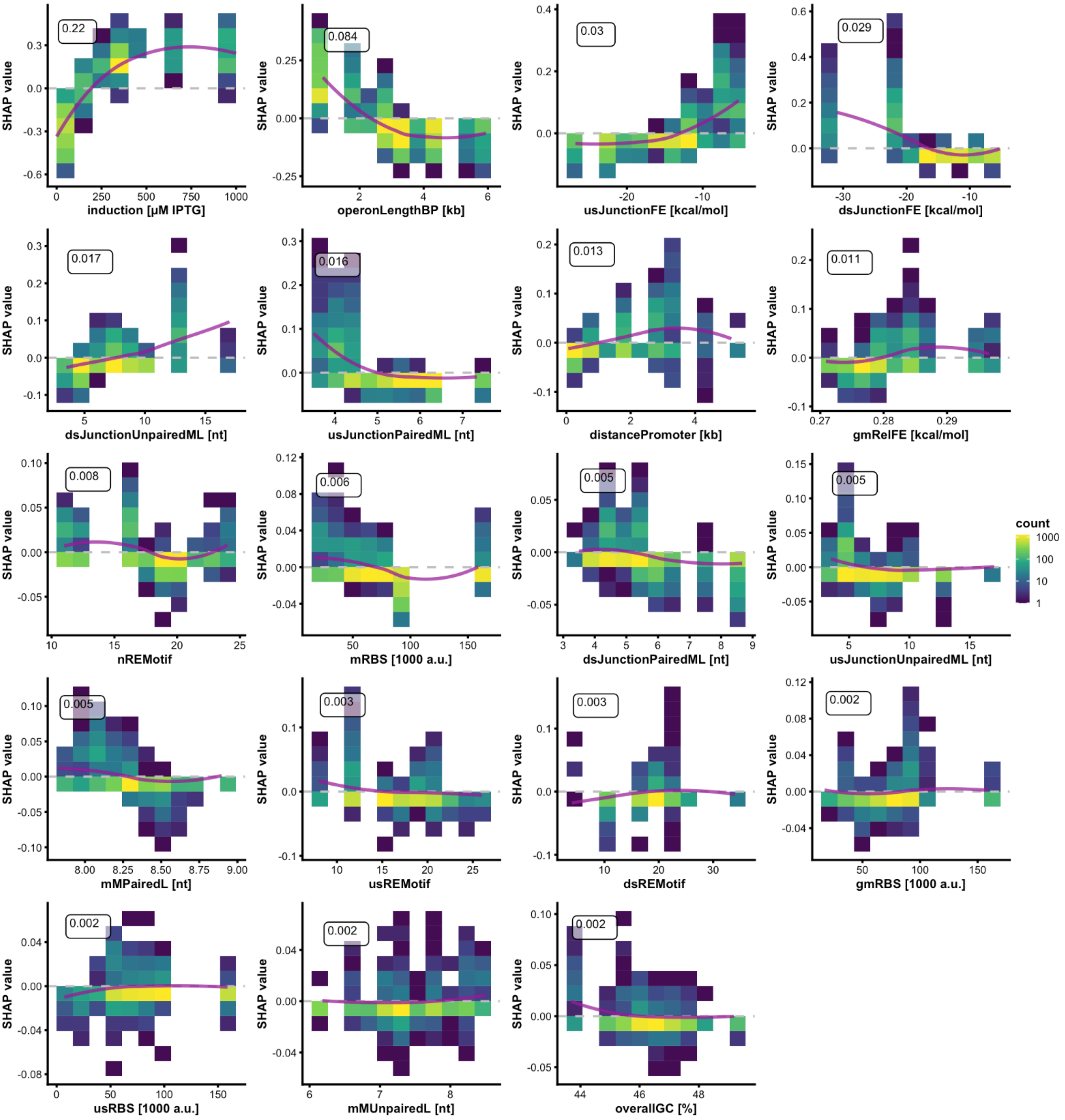


**Figure S8. SHAP value analysis for all features.** SHAP values plotted against feature values. The dashed horizontal line marks the position of points with a SHAP value of zero. The colored squares indicate how many values there are in within the indicated area of SHAP values and feature values. The purple line illustrates rough trends using a LOESS fit over the plotted data. The number at the top left of each plot is the mean absolute SHAP value of the corresponding feature. A detailed description corresponding to the feature abbreviations is given in **Table S5**.

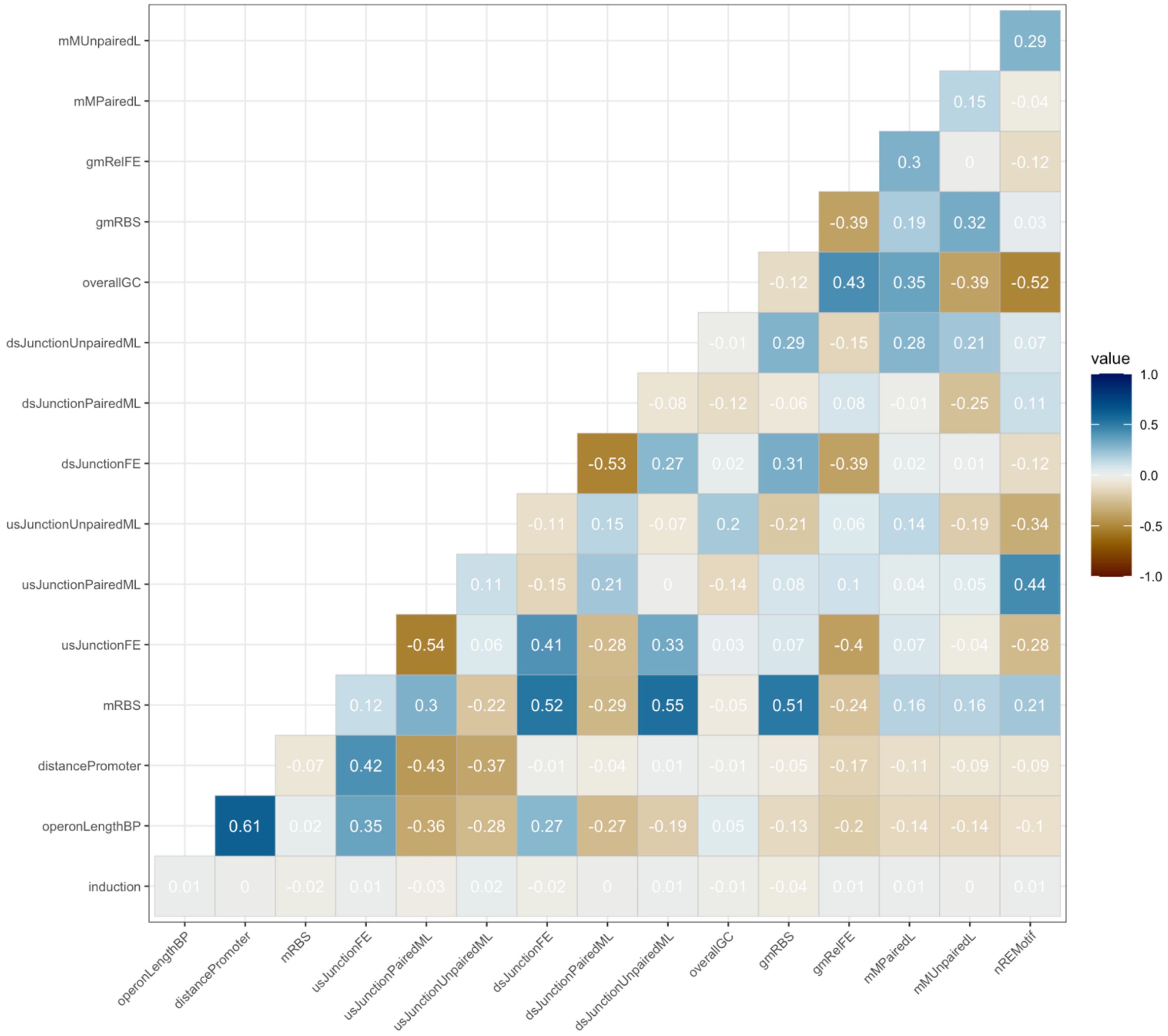


**Figure S9. Correlation analysis of all features.** The depicted correlation matrix contains the correlation coefficients for all features based on the complete data set.

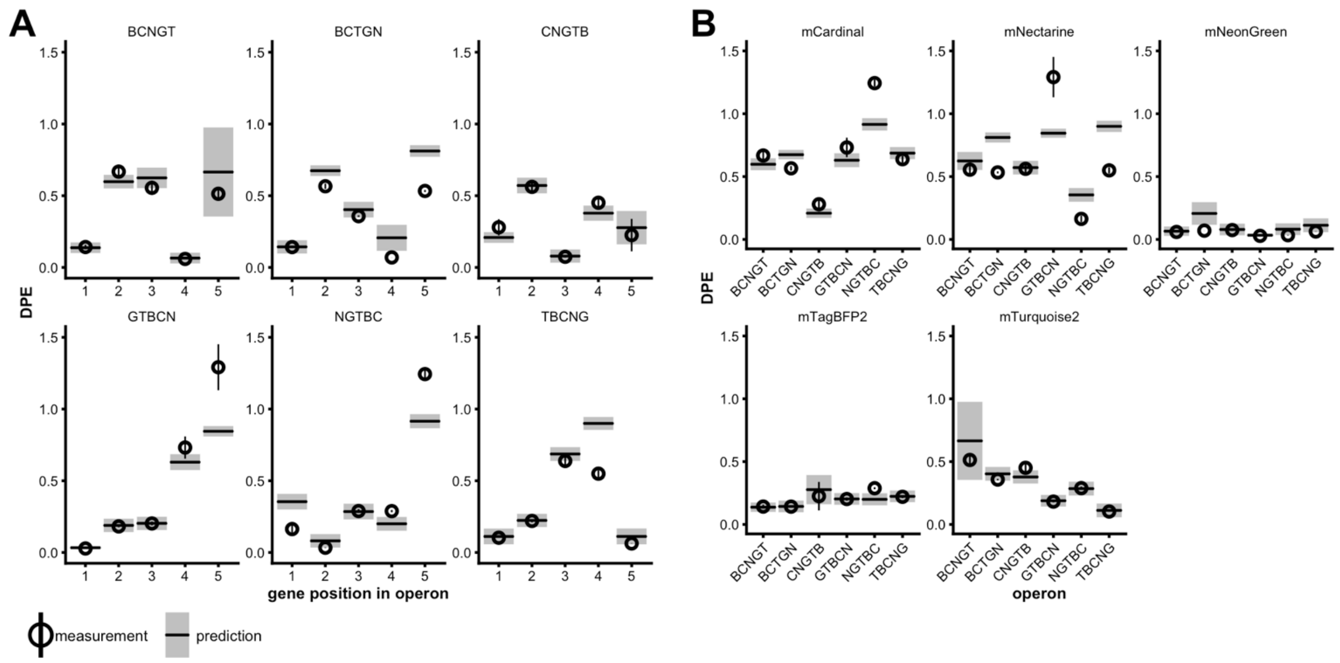


**Figure S10. Example operon predictions for operon permutations of five FP genes.** The measured DPE and the predicted DPE for the operon designs at an inducer concentration of 400 µM IPTG are plotted as circle with error bar and as horizontal line with grey area, respectively. The circles indicate the mean value of at least three biological replicates for each measurement, and the black horizontal lines indicate the corresponding single predicted value. The error bar range indicates the 95% confidence interval calculated using the standard error of the mean for measurements, and the grey area indicates the 95% confidence interval computed using the infinitesimal jackknife estimate of the standard error for predictions[43]. (**A**) Each panel represents one operon design indicated by the order of the letters above each panel (C: mCardinal, N: mNectarine, G: mNeonGreen, S: mT-Sapphire, T: mTurquoise2, B: mTagBFP2). (B) Each panel shows the different measured and predicted DPE values of one of the FPs that are part of the different operon designs (horizontal axis).

**Table S1. List of DNA fragments used in this project.** The here listed 20 fragments were amplified using the corresponding amplification primers (see **Table S2**) and subsequently assembled in LCRs using the appropriate bridging oligos (see **Table S3**) to form all plasmids listed in **Table S4**.

| Fragment ID | Description | Forward amplification primer ID | Reverse amplification primer ID | Sequence (5' → 3') |
|---|---|---|---|---|
| fDG137 | mCardinal | oDG306 | oDG307 | TAGTAAATATAATCGTATAGAATAGGAGGAACCATGGTCAGCAAAGGAGAGGAATT<br>AATCAAAGAAAATATGCATATGAAGCTGTATATGGAGGGTACGGTGAATAACCATC<br>ACTTTAAGTGTACCACCGAGGGGGAGGGAAAACCCTATGAGGGGACCCAAACCCAG<br>CGGATTAAGGTCGTAGAAGGCGGACCGCTGCCGTTCGCATTTGACATTCTGGCTAC<br>ATGTTTTATGTACGGCAGCAAAACCTTTATCAATCATACCCAGGGCATTCCTGATT<br>TCTTTAAACAGTCCTTCCCAGAGGGTTTTACCTGGGAGCGTGTCACCACCTACGAA<br>GACGGTGGTGTTCTGACCGTTACACAAGATACGAGTTTGCAGGACGGTTGCCTGAT<br>TTATAACGTTAAACTGCGTGGCGTGAATTTTCCGAGCAACGGGCCCGTAATGCAGA<br>AAAAGACGCTGGGCTGGGAAGCAACCACTGAAACCTTGTATCCAGCAGACGGTGGC<br>CTGGAAGGTCGTTGTGATATGGCACTCAAACTGGTTGGAGGCGGTCATTTACATTG<br>CAATCTGAAGACAACTTACCGTTCCAAAAAACCGGCCAAAAACCTGAAAATGCCTG<br>GCGTTTATTTTGTTGATCGTCGTCTGGAACGCATTAAAGAAGCAGATAACGAAACC<br>TACGTAGAACAACATGAAGTCGCGGTGGCGCGCTATTGCGATCTGCCCAGCAAACT<br>TGGTCATAAATTAAACGGTATGGACGAACTCTATAAACATCATCACCACCATCACG<br>AAGGTAAAAGCAGCGGTAGCGGTAGTGAAAGCAAAAGCACCGATTACAAAGACGAC<br>GACGACAAATAATAA |
| fDG138 | mNectarine | oDG308 | oDG309 | CGAGACGATACAATAGAAGGAGGAGGCACACAAATGGTTAGCAAAGGCGAAGAAGA<br>TAACATGGCCATTATCAAGGAGTTTATGCGTTTCAAAGTTCATATGGAGGGTTCCG<br>TCAACGGGCATGAGTTCGAGATTGAAGGCGAGGGGGAGGGCCGTCCGTACGAGGGT<br>ACACAAACAGCCAAACTGAAAGTCACGAAGGGTGGACCACTTCCGTTCGCGTGGGA<br>TATCCTGTCACCTCAATTCTGCTATGGAAGCAAAGCGTACGTGAAACATCCGGCCG<br>ATATTCCGGACTATCTGAAACTGTCGTTCCCTGAAGGTTTAAACTGGGAACGTGTG<br>ATGAACTTTGAGGACGGCGGGGTTGTTACCGTAACGCAGGATTCCTCTCTGCAAGA<br>CGGCGAGTTCATTTACAAGGTCAAACTCCGTGGTACAAATTTTCCCAGTGATGGTC<br>CAGTTATGCAGTGTCGCACCGTTGGTTGGGAGGCCAGTACCGAACGTATGCATCCG<br>GAGGACGGCGCACTCAAAGGCGAAATCATGCAACGCTTAAAGCTCAAGGACGGCGG<br>TCATTACGACGCGGAAGTCAAAACAACTTACAAAGCAAAAAAACCTGTGCAGTTAC<br>CGGGTGCATATAACGTCGACATTAAACTCGATATTCTTTCCCATAACGAGGACTAC<br>ACAATCGTTGAGCTGTACGAACGCGCGGAAGGCCGTCACAGCACGGGTGGATGGA<br>CGAGCTCTATAAGCATCATCACCACCATCACGAAGGAAAATCGTCCGGGAGTGGGA<br>GCGAAAGTAAATCAACTGAACAAAAACTGATTAGCGAAGAAGACCTGTAATAA |
| fDG139 | LSSmOrange | oDG310 | oDG311 | AGATCGGACAAGGATCTAAGGAAGAGAGGTTAAGAATGGTGAGCAAAGGCGAAGAA<br>AATAACATGGCCATCATCAAAGAATTTATGCGCTTTAAAGTGCGCATGGAAGGTAG<br>CGTTAACGGCCACGAATTTGAAATTGAAGGAGAAGGCGAAGGTCGTCCGTACGAAG<br>GTTTTCAGACCGTTAAACTGAAAGTGACCAAAGGTGGTCCGCTCCCGTTTGCATGG<br>GATATTCTGAGTCCGCAGTTTACCTATGGTAGCAAAGCATATGTGAAACACCCGGC<br>AGATATCCCGGATTATCTGAAACTGAGCTTTCCGGAAGGTTTTAAGTGGGAACGTG<br>TCATGAATTTTGAAGATGGTGGTGTTGTTACCGTTACCCAGGATAGCAGCCTGCAA<br>GATGGTGAATTTATCTACAAAGTTAAACTGCGTGGCACCAACTTTCCGAGTGATGG<br>TCCGGTTATGCAGAAAAAAACCATGGGTATGGAAGCAAGCAGCGAACGTATGTATC<br>CGGAAGATGGCGCACTGAAAGGTGAAGATAAACTGCGCCTGAAACTGAAAGACGGT<br>GGCCACTATACCAGCGAAGTTAAAACCACCTACAAAGCCAAAAAACCTGTTCAACT<br>GCCTGGTGCCTATATCGTTGATATTAAACTGGATATCACCAGCCACAACGAGGATT<br>ATACCATCGTCGAACAGTATGAACGTGCAGAAGGTCGCCACAGCACAGGTGGTATG<br>GACGAACTGTATAAACATCATCACCACCATCACGAAGGTAAGTCGTCAGGTAGCGG<br>ATCGGAATCAAAGTCTACCGGTAAACCGATTCCGAATCCGCTGCTGGGTCTGGATA<br>GCACCTAATAA |
| fDG140 | mNeonGreen | oDG312 | oDG313 | GAAAGAAAACAGCACTCCATAGAAGGAGGTTTATATGGTCAGCAAAGGTGAAGAAG<br>ATAATATGGCAAGCCTCCCAGCAACCCACGAACTGCACATTTTTGGTAGCATTAAC<br>GGCGTGGATTTTGATATGGTTGGTCAGGGTACAGGTAATCCGAACGACGGTTATGA<br>AGAACTGAATCTGAAAAGCACCAAAGGCGATCTGCAATTTAGCCCGTGGATTCTGG<br>TTCCGCACATTGGTTATGGTTTTCACCAGTATCTGCCGTATCCGGATGGTATGAGC<br>CCGTTTCAAGCAGCAATGGTTGATGGTAGCGGTTATCAGGTTCACCGTACCATGCA<br>GTTTGAAGACGGTGCAAGCCTGACCGTTAATTATCGTTATACCTATGAAGGCAGCC<br>ACATTAAAGGCGAAGCACAGGTTAAAGGTACAGGTTTTCCGGCAGATGGTCCGGTT<br>ATGACCAATAGTCTGACCGCAGCAGATTGGTGTCGTAGCAAAAAAACCTATCCGAA<br>CGATAAAACCATCATCAGCACCTTCAAATGGTCATATACCACCGGCAATGGTAAAC<br>GTTATCGTAGCACCGCACGTACCACCTATACCTTTGCAAAACCGATGGCAGCAAAC<br>TATCTGAAAAATCAGCCGATGTATGTGTTCCGTAAAACCGAACTGAAACACAGCAA<br>AACAGAGCTGAACTTTAAAGAGTGGCAGAAAGCCTTTACCGATGTGATGGGTATGG<br>ACGAGCTCTACAAACATCATCACCACCATCACGAAGGGAAAAGCTCTGGATCGGGT<br>TCCGAAAGCAAATCGACCTATCCGTATGATGTTCCGGATTATGCATAATAA |

| | | | | |
|---|---|---|---|---|
| fDG141 | mT-Sapphire | oDG314 | oDG315 | ATTATAGGTCAAAAAATAGAGGAGGACGGAAATCAATGGTCAGCAAAGGTGAAGAA<br>CTGTTTACAGGTGTTGTTCCGATTCTGGTTGAACTGGATGGTGACGTTAACGGCCA<br>CAAATTTTCAGTTAGCGGTGAAGGCGAAGGTGATGCAACCTATGGTAAACTGACCC<br>TGAAATTTATCTGTACCACCGGCAAACTGCCGGTTCCGTGGCCGACCCTGGTTACC<br>ACCTTTAGCTATGGTGTTATGGTTTTTGCACGTTATCCGGATCATATGAAACAGCA<br>CGATTTTTTCAAAAGCGCAATGCCGGAAGGTTACGTTCAAGAACGTACCATCTTCT<br>TCAAAGATGACGGCAACTATAAAACCCGTGCCGAAGTTAAATTTGAAGGTGATACC<br>CTGGTGAATCGCATTGAACTGAAAGGCATCGATTTTAAAGAGGATGGTAATATCCT<br>CGGCCACAAACTGGAATATAATTTCAATAGCCACAACGTGTATATCATGGCCGACA<br>AACAGAAAAATGGCATCAAAGCCAATTTCAAAATCCGCCACAATATTGAAGATGGT<br>GGTGTTCAACTGGCAGATCACTATCAGCAGAATACCCCGATTGGTGATGGTCCGGT<br>TCTGCTGCCGGATAATCACTATCTGAGCATTCAGAGCAAACTGAGCAAAGATCCGA<br>ACGAAAAACGTGATCACATGGTGCTGCTGGAATTTGTTACCGCAGCAGGTATTACC<br>CTGGGTATGGATCAACTCTATAAACATCATCACCACCATCACGAGGGAAAGTCTAG<br>CGGATCTGGAAGTGAGTCTAAATCAACAAATTGGAGCCATCCGCAGTTTGAAAAAT<br>AATAA |
| fDG142 | mTurquoise2 | oDG316 | oDG317 | AGGCAGTTCGCCAGTAAGTAAGGAGGTAATCAATGGTATCCAAGGGCGAAGAACTC<br>TTCACCGGGGTTGTCCCTATCCTGGTTGAACTGGATGGTGACGTCAATGGTCACAA<br>ATTTTCGGTGAGCGGCGAGGGCGAGGGTGATGCCACGTACGGAAAGCTCACACTCA<br>AGTTTATTTGTACAACCGGGAAATTACCTGTTCCGTGGCCAACTCTCGTAACAACA<br>CTGTCCTGGGGCGTCCAGTGTTTCGCTCGCTATCCAGACCATATGAAACAGCACGA<br>CTTCTTTAAAAGCGCTATGCCGGAAGGTTACGTTCAGGAACGTACCATTTTCTTTA<br>AAGATGACGGCAATTACAAAACCCGTGCTGAGGTAAAGTTCGAAGGGGACACTCTG<br>GTGAATCGCATTGAGTTGAAGGGCATCGACTTTAAAGAAGACGGTAACATTCTCGG<br>TCACAAACTCGAATACAATTACTTTAGCGACAACGTGTATATCACCGCGGATAAGC<br>AGAAAAACGGCATTAAGGCAAATTTTAAAATCCGTCACAACATTGAAGACGGGGGA<br>GTACAATTGGCCGACCACTATCAGCAAAACACGCCAATTGGTGACGGTCCGGTTCT<br>GTTACCCGATAACCATTACCTGTCTACCCAGTCAAAACTGTCGAAAGACCCAAACG<br>AAAAACGGGACCACATGGTACTGTTGGAGTTTGTTACTGCGGCGGGCATCACCTTG<br>GGCATGGACGAGCTCTATAAACATCATCACCACCATCACGAGGGGAAGTCAAGTGG<br>GTCTGGGTCTGAAAGTAAATCAACCATGGCGAGCATGACTGGTGGACAGCAAATGG<br>GTTAATAA |
| fDG143 | mTagBFP2 | oDG318 | oDG319 | TATTTTAATAGGAGTCAGTAAGGAGAATACGGATGGTCAGCAAAGGTGAAGAACTC<br>ATCAAAGAAAACATGCACATGAAACTGTACATGGAAGGCACCGTCGATAACCACCA<br>CTTTAAATGTACCAGCGAAGGTGAAGGTAAACCGTACGAAGGCACCCAGACCATGC<br>GTATTAAAGTTGTTGAAGGTGGTCCGCTGCCGTTTGCATTTGATATTCTGGCAACC<br>AGCTTTCTGTATGGTAGCAAAACCTTTATTAACCACACCCAGGGTATCCCGGATTT<br>TTTCAAACAGAGCTTTCCGGAAGGTTTTACCTGGGAACGTGTTACCACCTATGAAG<br>ATGGTGGTGTTCTGACCGCAACCCAGGATACCAGCCTGCAAGATGGTTGTCTGATT<br>TATAATGTGAAAATCCGTGGCGTGAACTTTACCAGCAATGGTCCGGTTATGCAGAA<br>AAAAACCCTGGGTTGGGAAGCATTTACCGAAACCCTGTATCCGGCAGATGGTGGCC<br>TGGAAGGTCGTAACGATATGGCACTGAAACTGGTTGGTGGTAGCCACCTGATTGCA<br>AATGCCAAAACCACCTATCGTAGCAAAAAACCGGCAAAAAATCTCAAAATGCCAGG<br>CGTGTATTATGTGGATTATCGTCTCGAACGCATTAAAGAAGCAAACAACGAAACCT<br>ATGTGGAACAGCACGAAGTTGCAGTTGCACGTTATTGTGATCTGCCGAGCAAACTG<br>GGTCACAAACTGAACCATCATCACCACCATCACGAAGGTAAGTCAAGCGGTTCTGG<br>ATCGGAAAGCAAGAGTACCTATACCGATATTGAAATGAACCGTCTGGGTAAATAAT<br>AA |
| fDG163 | pSEVA2xx<br>(kanR) | oDG304 | oDG414 | ACTAGTCTTGGACTCCTGTTGATAGATCCAGTAATGACCTCAGAACTCCATCTGGA<br>TTTGTTCAGAACGCTCGGTTGCCGCCGGGCGTTTTTTATTGGTGAGAATCCAGGGG<br>TCCCCAATAATTACGATTTAAATTTGTGTCTCAAAATCTCTGATGTTACATTGCAC<br>AAGATAAAAATATATCATCATGAACAATAAAACTGTCTGCTTACATAAACAGTAAT<br>ACAAGGGGTGTTATGAGCCATATTCAGCGTGAAACGAGCTGTAGCCGTCCGCGTCT<br>GAACAGCAACATGGATGCGGATCTGTATGGCTATAAATGGGCGCGTGATAACGTGG<br>GTCAGAGCGGCGCGACCATTTATCGTCTGTATGGCAAACCGGATGCGCCGGAACTG<br>TTTCTGAAACATGGCAAAGGCAGCGTGGCGAACGATGTGACCGATGAAATGGTGCG<br>TCTGAACTGGCTGACCGAATTTATGCCGCTGCCGACCATTAAACATTTTATTCGCA<br>CCCCGGATGATGCGTGGCTGCTGACCACCGCGATTCCGGGCAAAACCGCGTTTCAG<br>GTGCTGGAAGAATATCCGGATAGCGGCGAAAACATTGTGGATGCGCTGGCCGTGTT<br>TCTGCGTCGTCTGCATAGCATTCCGGTGTGCAACTGCCCGTTTAACAGCGATCGTG<br>TGTTTCGTCTGGCCCAGGCGCAGAGCCGTATGAACAACGGCCTGGTGGATGCGAGC<br>GATTTTGATGATGAACGTAACGGCTGGCCGGTGGAACAGGTGTGGAAAGAAATGCA<br>TAAACTGCTGCCGTTTAGCCCGGATAGCGTGGTGACCCACGGCGATTTTAGCCTGG<br>ATAACCTGATTTTCGATGAAGGCAAACTGATTGGCTGCATTGATGTGGGCCGTGTG<br>GGCATTGCGGATCGTTATCAGGATCTGGCCATTCTGTGGAACTGCCTGGGCGAATT<br>TAGCCCGAGCCTGCAAAAACGTCTGTTTCAGAAATATGGCATTGATAATCCGGATA<br>TGAACAAACTGCAATTTCATCTGATGCTGGATGAATTTTTCTAATAATTAATTGGA<br>CAAGGGTCCTTTTCCGCTGCATAACCCTGCTTCGGGGTCATTATAGCG |

| | | | | |
|---|---|---|---|---|
| fDG164 | pSEVAx9x (pBR322-ROP ori) | oDG382 | oDG413 | ATTTTTTCGGTATATCCATCCTTTTTCGCACGATATACAGGATTTTGCCAAAGGGT<br>TCGTGTAGACTTTCCTTGGTGTATCCAACGGCGTCAGCCGGGCAGGATAGGTGAAG<br>TAGGCCCACCCGCGAGCGGGTGTTCCTTCTTCACTGTCCCTTATTCGCACCTGGCG<br>GTGCTCAACGGGAATCCTGCTCTGCGAGGCTGGCCGTAGGCCGGCCCCGTAGAAAA<br>GATCAAAGGATCTTCTTGAGATCCTTTTTTTCTGCGCGTAATCTGCTGCTTGCAAA<br>CAAAAAAACCACCGCTACCAGCGGTGGTTTGTTTGCCGGATCAAGAGCTACCAACT<br>CTTTTTCCGAAGGTAACTGGCTTCAGCAGAGCGCAGATACCAAATACTGTCCTTCT<br>AGTGTAGCCGTAGTTAGGCCACCACTTCAAGAACTCTGTAGCACCGCCTACATACC<br>TCGCTCTGCTAATCCTGTTACCAGTGGCTGCTGCCAGTGGCGATAAGTCGTGTCTT<br>ACCGGGTTGGACTCAAGACGATAGTTACCGGATAAGGCGCAGCGGTCGGGCTGAAC<br>GGGGGGTTCGTGCACACAGCCCAGCTTGGAGCGAACGACCTACACCGAACTGAGAT<br>ACCTACAGCGTGAGCTATGAGAAAGCGCCACGCTTCCCGAAGGGAGAAAGGCGGAC<br>AGGTATCCGGTAAGCGGCAGGGTCGGAACAGGAGAGCGCACGAGGGAGCTTCCAGG<br>GGGAAACGCCTGGTATCTTTATAGTCCTGTCGGGTTTCGCCACCTCTGACTTGAGC<br>GTCGATTTTTGTGATGCTCGTCAGGGGGGCGGAGCCTATGGAAAAACGCCAGCAAC<br>GCGGCCTTTTTACGGTTCCTGGCCTTTTGCTGGCCTTTTGCTCACATGTTCTTTCC<br>TGCGTTATCCCCTGATTCTGTGGATAACCGTATTACCGCCTTTGAGTGAGCTGATA<br>CCGCTCGCCGCAGCCGAACGACCGAGCGCAGCGAGTCAGTGAGCGAGGAAGCGGAA<br>GAGCGCCTGATGCGGTATTTTCTCCTTACGCATCTGTGCGGTATTTCACACCGCAT<br>ATGGTGCACTCTCAGTACAATCTGCTCTGATGCCGCATAGTTAAGCCAGTATACAC<br>TCCGCTATCGCTACGTGACTGGGTCATGGCTGCGCCCCGACACCCGCCAACACCCG<br>CTGACGCGCCCTGACGGGCTTGTCTGCTCCCGGCATCCGCTTACAGACAAGCTGTG<br>ACCGTCTCCGGGAGCTGCATGTGTCAGAGGTTTTCACCGTCATCACCGAAACGCGC<br>GAGGCAGCTGCGGTAAAGCTCATCAGCGTGGTCGTGCAGCGATTCACAGATGTCTG<br>CCTGTTCATCCGCGTCCAGCTCGTTGAGTTTCTCCAGAAGCGTTAATGTCTGGCTT<br>CTGATAAAGCGGGCCATGTTAAGGGCGGTTTTTTCCTGTTTGGTCACTGATGCCTC<br>CGTGTAAGGGGGATTTCTGTTCATGGGGGTAATGATACCGATGAAACGAGAGAGGA<br>TGCTCACGATACGGGTTACTGATGATGAACATGCCCGGTTACTGGAACGTTGTGAG<br>GGTAAACAACTGGCGGTATGGATGCGGCGGGGGCGCGCCCAGCTGTCTAGGGCGGC<br>GGATTTGTCCTACTCAGGAGAGCGTTCACCGACAAACAACAGATAAAACGAAAGGC<br>CCAGTCTTTCGACTGAGCCTTTCGTTTTATTTGATGCCTTTAATTAA |
| fDG197 | lacI_pT7lacO | oDG305 | oDG488 | CTATCACTGCCCGCTTTCCAGTCGGGAAACCTGTCGTGCCAGCTGCATTAATGAAT<br>CGGCCAACGCGCGGGGAGAGGCGGTTTGCGTATTGGGCGCCAGGGTGGTTTTTCTT<br>TTCACCAGTGAGACGGGCAACAGCTGATTGCCCTTCACCGCCTGGCCCTGAGAGAG<br>TTGCAGCAAGCGGTCCACGCTGGTTTGCCCCAGCAGGCGAAAATCCTGTTTGATGG<br>TGGTTAACGGCGGGATATAACATGAGCTGTCTTCGGTATCGTCGTATCCCACTACC<br>GAGATATCCGCACCAACGCGCAGCCCGGACTCGGTAATGGCGCGCATTGCGCCCAG<br>CGCCATCTGATCGTTGGCAACCAGCATCGCAGTGGGAACGATGCCCTCATTCAGCA<br>TTTGCATGGTTTGTTGAAAACCGGACATGGCACTCCAGTCGCCTTCCCGTTCCGCT<br>ATCGGCTGAATTTGATTGCGAGTGAGATATTTATGCCAGCCAGCCAGACGCAGACG<br>CGCCGAGACAGAACTTAATGGGCCCGCTAACAGCGCGATTTGCTGGTGACCCAATG<br>CGACCAGATGCTCCACGCCCAGTCGCGTACCGTCTTCATGGGAGAAAATAATACTG<br>TTGATGGGTGTCTGGTCAGAGACATCAAGAAATAACGCCGGAACATTAGTGCAGGC<br>AGCTTCCACAGCAATGGCATCCTGGTCATCCAGCGGATAGTTAATGATCAGCCCAC<br>TGACGCGTTGCGCGAGAAGATTGTGCACCGCCGCTTTACAGGCTTCGACGCCGCTT<br>CGTTCTACCATCGACACCACCACGCTGGCACCCAGTTGATCGGCGCGAGATTTAAT<br>CGCCGCGACAATTTGCGACGGCGCGTGCAGGGCCAGACTGGAGGTGGCAACGCCAA<br>TCAGCAACGACTGTTTGCCCGCCAGTTGTTGTGCCACGCGGTTGGGAATGTAATTC<br>AGCTCCGCCATCGCCGCTTCCACTTTTTCCCGCGTTTTCGCAGAAACGTGGCTGGC<br>CTGGTTCACCACGCGGGAAACGGTCTGATAAGAGACACCGGCATACTCTGCGACAT<br>CGTATAACGTTACTGGTTTCACATTCACCACCCTGAATTGACTCTCTTCCGGGCGC<br>TATCATGCCATACCGCGAAAGGTTTTGCGCCATTCGATGGTGTCAGGTTCTGTTAA<br>GTAACTGAACCCAATGTCGTTAGTGACGCTTACCTCTTAAGAGGTCACTGACCTAA<br>CATAATACGACTCACTATAGGGGAATTGTGAGCGGATAACAATTCC |
| fDG201 | XylE | oDG483 | oDG315 | GCTCCTAGCATATAAGGAGAATTTTTATGAACAAAGGTGTTATGCGTCCGGGTCAC<br>GTTCAACTGCGTGTTCTGGATATGAGCAAAGCACTGGAACACTACGTTGAACTGCT<br>GGGTCTGATTGAAATGGATCGTGACGATCAGGGTCGTGTTTATCTGAAAGCCTGGA<br>CCGAAGTTGATAAATTCAGCCTGGTTCTGCGTGAAGCAGACGAACCTGGTATGGAT<br>TTTATGGGTTTTAAAGTTGTTGACGAGGACGCACTGCGTCAACTGGAACGTGATCT<br>GATGGCCTACGGTTGTGCCGTTGAACAGTTACCGGCAGGCGAACTGAATAGCTGTG<br>GTCGTCGTGTTCGTTTTCAGGCACCGAGCGGTCACCACTTTGAACTGTACGCAGAT<br>AAAGAATATACCGGCAAGTGGGGTCTGAACGACGTTAATCCGGAAGCCTGGCCTCG<br>CGATCTGAAAGGTATGGCAGCAGTTCGTTTTGATCACGCACTGATGTACGGCGACG<br>AACTGCCTGCAACCTACGACCTGTTTACCAAAGTTCTGGGTTTTTATCTGGCAGAA<br>CAGGTTCTGGACGAAAACGGCACCCGTGTTGCACAGTTTCTGAGCCTGAGCACCAA<br>AGCACACGACGTTGCCTTTATTCACCACCCGGAAAAAGGTCGTCTGCACCACGTTA<br>GCTTTCACCTGGAAACCTGGGAAGATTTACTGCGTGCAGCCGATCTGATTAGTATG<br>ACCGATACCAGTATTGATATTGGTCCGACACGTCACGGTCTGACCCACGGTAAAAC<br>AATCTATTTCTTTGATCCGAGCGGTAATCGCAACGAAGTTTTTTGTGGTGGCGATT<br>ATAACTATCCGGATCACAAACCTGTTACCTGGACCACCGATCAACTGGGTAAAGCA<br>ATCTTTTATCACGATCGTATTCTGAACGAACGCTTTATGACCGTTCTGACCCATCA<br>TCACCACCATCACGAGGGAAAGTCTAGCGGATCTGGAAGTGAGTCTAAATCAACAA<br>ATTGGAGCCATCCGCAGTTTGAAAAATAATAA |

| | | | | |
|---|---|---|---|---|
| fDG202 | DocS | oDG484 | oDG485 | TAAATTCTTATCATTAAAATAAGGAGAGTCTCTATGCATCATCACCACCATCACGT<br>TGTTCCGGGTACACCGAGCACCAAACTGTACGGTGACGTTAACGACGACGGTAAAG<br>TTAATAGCACCGACGCAGTTGCACTGAAACGTTACGTTCTGCGTAGCGGTATTTCA<br>ATTAATACCGATAACGCCGATCTGAACGAGGACGGTCGTGTTAATTCAACCGATCT<br>GGGTATTCTGAAACGCTATATCCTGAAAGAAATTGATACCCTGCCGTATAAAAACT<br>AATAA |
| fDG210 | AspA | oDG536 | oDG537 | ACTGAAATCACAGACTCAACTCATTTAAGCGCAGGTTTTTTTATGCATCATCACCA<br>CCATCACGAAGGTAAAAGCAGCGGTAGCGGTAGTGAAAGCAAAAGCACCGATTACA<br>AAGACGACGACGACAAAAGCAACAATATTCGTATTGAAGAGGATCTGCTGGGCACC<br>CGTGAAGTTCCGGCAGACGCCTATTACGGTGTTCACACCCTGCGTGCAATTGAAAA<br>TTTCTATATCAGCAACAACAAAATCAGCGATATCCCGGAATTTGTTCGTGGTATGG<br>TTATGGTTAAAAAAGCAGCAGCAATGGCCAACAAAGAACTGCAAACAATTCCGAAA<br>AGCGTTGCCAACGCAATTATTGCAGCCTGTGACGAAGTTCTGAATAACGGTAAGTG<br>TATGGATCAGTTTCCGGTTGACGTTTATCAAGGTGGTGCAGGCACCAGCGTTAATA<br>TGAATACCAACGAAGTGCTGGCAAATATTGGTCTGGAACTGATGGGTCACCAGAAA<br>GGTGAATATCAGTATCTGAATCCGAACGATCACGTTAATAAGTGTCAGAGCACCAA<br>CGACGCCTATCCGACCGGCTTTCGTATTGCAGTTTATAGCAGCCTGATTAAACTGG<br>TGGACGCAATTAATCAACTGCGCGAAGGTTTTGAACGTAAAGCAGTTGAATTTCAG<br>GATATTCTGAAGATGGGTCGTACCCAGTTACAGGACGCAGTTCCGATGACACTGGG<br>TCAAGAATTTCGTGCCTTTAGTATTCTGCTGAAAGAAGAAGTGAAAAATATCCAGC<br>GTACCGCAGAACTGCTGCTGGAAGTTAATCTGGGTGCAACCGCAATTGGTACAGGT<br>CTGAATACCCCGAAAGAATATAGTCCGCTGGCAGTTAAAAAACTGGCCGAAGTTAC<br>AGGTTTTCCGTGTGTTCCTGCCGAGGATCTGATTGAAGCAACCAGCGATTGTGGTG<br>CCTACGTTATGGTTCACGGTGCACTGAAACGTCTGGCCGTTAAAATGAGCAAAATT<br>TGTAACGATCTGCGTCTGCTGAGCAGCGGTCCGCGTGCCGGTCTGAACGAAATTAA<br>CCTGCCGGAATTACAGGCAGGTAGCAGTATTATGCCTGCAAAAGTTAATCCGGTTG<br>TTCCGGAAGTTGTTAATCAGGTTTGCTTTAAAGTGATCGGTAACGATACCACCGTT<br>ACAATGGCAGCAGAAGCAGGTCAACTGCAACTGAACGTTATGGAACCTGTTATTGG<br>TCAGGCAATGTTTGAAAGCGTTCATATTCTGACCAACGCCTGTTATAATCTGCTGG<br>AAAAGTGTATTAACGGTATCACCGCCAATAAAGAAGTTTGCGAAGGTTACGTGTAT<br>AACAGTATTGGTATTGTGACCTATCTGAACCCGTTTATTGGCCACCACAACGGTGA<br>TATTGTTGGTAAAATTTGTGCAGAAACCGGCAAAAGCGTTCGTGAAGTTGTTCTGG<br>AACGTGGTCTGCTGACCGAAGCAGAACTGGACGATATCTTTAGCGTTCAGAATCTG<br>ATGCACCCTGCCTATAAAGCAAAACGTTATACCGACGAAAGCGAACAGTGATAA |
| fDG211 | PanD | oDG538 | oDG309 | ACTTCGTCAATCGTACTCATAAGGAGGTATATATGCTGCGTACAATTCTGGGTAGC<br>AAAATTCACCGTGCAACCGTTACACAGGCCGATCTGGATTACGTTGGTAGCGTTAC<br>AATTGACGCGGATCTGGTTCACGCAGCAGGTCTGATTGAAGGTGAAAAAGTTGCGA<br>TTGTGGATATTACCAACGGTGCACGTCTGGAAACCTACGTTATTGTTGGTGACGCA<br>GGTACAGGCAATATTTGTATTAACGGTGCAGCAGCACACCTGATTAATCCGGGTGA<br>TCTGGTGATTATTATGAGCTATCTGCAAGCAACCGACGCCGAAGCAAAAGCCTACG<br>AACCGAAAATTGTTCACGTTGACGCAGATAATCGTATTGTTGCACTGGGTAACGAT<br>CTGGCAGAAGCAATTCCTGGTAGCGGTCTGCTGACCAGCCGTAGTATTCATCATCA<br>CCACCATCACGAAGGAAAATCGTCCGGGAGTGGGAGCGAAAGTAAATCAACTGAAC<br>AAAAACTGATTAGCGAAGAAGACCTGTAATAA |
| fDG212 | BauA | oDG539 | oDG540 | ACTCCAGACATTTTTAAGGCAGGTTTTTATGCATCATCACCACCATCACGAAGGTA<br>AGTCGTCAGGTAGCGGATCGGAATCAAAGTCTACCGGTAAACCGATTCCGAATCCG<br>CTGCTGGGTCTGGATAGCACCAATCAGCCGCTGAACGTTGCACCGCCTGTTAGCAG<br>CGAACTGAATCTGCGTGCACACTGGATGCCGTTTAGCGCAAATCGTAATTTTCAGA<br>AAGATCCGCGTATTATTGTTGCAGCAGAAGGTAGCTGGCTGACCGACGATAAAGGT<br>CGTAAAGTGTACGATAGCCTGAGCGGTCTGTGGACCTGTGGTGCAGGTCACAGCCG<br>TAAAGAAATTCAAGAAGCAGTTGCCCGTCAACTGGGCACCCTGGATTATTCACCGG<br>GTTTTCAGTACGGTCACCCGCTGAGCTTTCAACTGGCAGAAAAAATTGCAGGTCTG<br>CTGCCTGGTGAACTGAACCACGTGTTTTTTACCGGCAGCGGTAGCGAGTGTGCAGA<br>TACCAGTATCAAAATGGCACGTGCGTATTGGCGTCTGAAAGGTCAGCCGCAGAAAA<br>CCAAACTGATTGGTCGTGCACGTGGTTATCACGGTGTTAACGTTGCAGGCACCAGC<br>TTAGGTGGTATTGGTGGTAATCGTAAAATGTTTGGTCAACTGATGGACGTTGATCA<br>CTTACCGCACACCTTACAGCCTGGTATGGCGTTTACCCGTGGTATGGCACAGACAG<br>GTGGTGTTGAACTGGCAAACGAACTGCTGAAACTGATCGAACTGCACGACGCAAGT<br>AATATTGCAGCAGTTATTGTTGAACCGATGAGCGGTAGTGCCGGTGTTCTGGTTCC<br>GCCTGTTGGTTATCTGCAACGTCTGCGTGAAATTTGTGATCAGCACAATATCCTGC<br>TGATCTTTGACGAAGTTATTACCGCCTTTGGTCGTCTGGGCACCTATAGCGGTGCA<br>GAATATTTTGGTGTTACACCGGATCTGATGAACGTGGCAAAACAGGTTACCAACGG<br>TGCAGTTCCGATGGGTGCCGTTATTGCAAGCAGCGAAATTTACGATACCTTTATGA<br>ATCAGGCACTGCCGGAACACGCAGTTGAATTTAGCCACGGTTATACCTATAGTGCA<br>CACCCGGTTGCCTGTGCAGCAGGTCTGGCAGCACTGGATATTCTGGCACGTGATAA<br>TCTGGTTCAGCAGAGCGCAGAACTGGCACCGCACTTTGAAAAAGGTCTGCACGGTC<br>TGCAAGGTGCAAAAAACGTTATTGATATTCGTAATTGTGGCCTGGCAGGCGCAATT<br>CAGATTGCACCGCGTGACGGTGATCCGACCGTTCGTCCGTTTGAAGCAGGTATGAA<br>ACTGTGGCAGCAGGGTTTTTACGTTCGTTTTGGTGGCGATACCTTACAGTTTGGTC<br>CGACCTTTAACGCACGTCCGGAAGAACTGGATCGTCTGTTTGACGCAGTTGGTGAA<br>GCACTGAACGGTATTGCCTAATAA |

| | | | | |
|---|---|---|---|---|
| fDG213 | YneI | oDG541 | oDG542 | ACAAGACATTTAATTAAGGACAGGTACGTTTATGCATCATCACCACCATCACGAAG<br>GGAAAAGCTCTGGATCGGGTTCCGAAAGCAAATCGACCTATCCGTATGATGTTCCG<br>GATTATGCAACAATTACACCGGCAACACACGCGATTAGTATTAATCCGGCAACAGG<br>TGAACAACTGAGCGTTCTGCCGTGGGCAGGCGCAGACGATATTGAAAACGCACTGC<br>AACTGGCAGCAGCAGGTTTTCGTGATTGGCGTGAAACCAATATTGATTATCGTGCA<br>GAAAAACTGCGCGATATTGGTAAAGCACTGCGTGCACGTAGCGAAGAAATGGCACA<br>GATGATTACCCGTGAAATGGGTAAACCGATTAATCAGGCACGTGCAGAAGTTGCAA<br>AAAGCGCAAATCTGTGTGATTGGTACGCAGAACACGGTCCGGCAATGCTGAAAGCA<br>GAACCGACACTGGTTGAAAATCAGCAGGCAGTTATTGAATATCGTCCGCTGGGTAC<br>AATTCTGGCAATTATGCCGTGGAATTTTCCGCTGTGGCAGGTTATGCGTGGTGCAG<br>TTCCGATTATTCTGGCAGGTAACGGTTATCTGCTGAAACACGCACCGAACGTTATG<br>GGTTGTGCCCAACTGATTGCACAGGTTTTTAAAGACGCAGGTATTCCGCAGGGTGT<br>TTACGGTTGGCTGAACGCAGATAACGACGGTGTTAGCCAGATGATCAAAGATAGCC<br>GTATTGCAGCAGTTACCGTTACAGGTAGCGTTCGTGCCGGTGCAGCAATTGGTGCC<br>CAAGCGGGTGCAGCACTGAAAAAGTGTGTTCTGGAATTAGGTGGTAGCGATCCGTT<br>TATTGTTCTGAACGACGCCGATCTGGAACTGGCAGTTAAAGCAGCAGTTGCAGGTC<br>GTTATCAGAATACAGGTCAGGTTTGTGCAGCAGCCAAACGCTTTATTATCGAAGAA<br>GGTATTGCAAGCGCCTTTACCGAACGCTTTGTTGCAGCCGCAGCAGCCCTGAAAAT<br>GGGTGATCCGCGTGACGAAGAAAACGCCCTGGGTCCGATGGCACGTTTTGATCTGC<br>GCGACGAGCTGCACCACCAGGTTGAAAAAACCCTGGCACAGGGTGCACGTCTGCTG<br>TTAGGTGGTGAAAAAATGGCTGGTGCAGGTAACTATTATCCGCCTACCGTTCTGGC<br>AAACGTTACACCGGAAATGACCGCCTTTCGTGAAGAAATGTTTGGTCCGGTTGCCG<br>CAATTACAATTGCAAAAGACGCCGAACACGCCTTAGAACTGGCCAACGATAGCGAA<br>TTTGGTCTGAGCGCAACAATTTTTACCACCGACGAAACCCAGGCACGCCAGATGGC<br>AGCACGTTTAGAGTGTGGTGGTGTGTTTATTAACGGCTATTGTGCAAGCGACGCAC<br>GTGTTGCCTTTGGCGGTGTGAAAAAAAGCGGTTTTGGTCGTGAACTGAGCCACTTT<br>GGTCTGCACGAATTTTGTAATATTCAGACCGTTTGGAAAGATCGTATCTAATAA |
| fDG214 | MatB | oDG543 | oDG544 | GAATTTAAGCGCTTAAGGAGAGTCATATATGCATCATCACCACCATCACGAGGGGA<br>AGTCAAGTGGGTCTGGGTCTGAAAGTAAATCAACCATGGCGAGCATGACTGGTGGA<br>CAGCAAATGGGTAGCAATCACCTGTTTGACGCAATGCGTGCAGCAGCACCGGGTAA<br>CGCACCGTTTATTCGTATTGATAATACCCGTACCTGGACCTACGACGACGCCTTTG<br>CACTGAGCGGTCGTATTGCAAGCGCAATGGACGCACTGGGTATTCGTCCGGGTGAT<br>CGTGTTGCAGTTCAGGTTGAAAAAAGCGCAGAAGCACTGATTCTGTATCTGGCCTG<br>TCTGCGTAGCGGTGCAGTTTATCTGCCGCTGAATACCGCGTATACCCTGGCAGAAC<br>TGGATTACTTTATTGGTGACGCAGAACCGCGTCTGGTTGTTGTTGCAAGCAGCGCA<br>CGTGCCGGTGTTGAAACAATTGCAAAACCGCGTGGTGCGATTGTTGAAACCCTGGA<br>CGCAGCAGGTAGCGGTAGCCTGCTGGATCTGGCACGTGACGAACCGGCAGATTTTG<br>TTGACGCAAGCCGTAGCGCAGACGATCTGGCAGCAATTCTGTATACCAGCGGCACC<br>ACAGGTCGTAGCAAAGGTGCAATGCTGACCCACGGTAATCTGCTGAGCAACGCACT<br>GACCCTGCGTGATTTTTGGCGTGTTACCGCAGGCGATCGTCTGATTCACGCACTGC<br>CGATTTTTCACACCCACGGTCTGTTTGTTGCAACCAACGTTACCCTGTTAGCCGGT<br>GCCTCAATGTTTCTGCTGAGTAAATTTGATCCGGAAGAAATTCTGAGCCTGATGCC<br>GCAGGCAACAATGCTGATGGGTGTTCCGACCTTTTACGTTCGTCTGCTGCAAAGCC<br>CTCGTCTGGATAAACAGGCAGTTGCAAATATTCGTCTGTTTATTAGCGGTAGTGCA<br>CCGCTGCTGGCAGAAACCCACACCGAATTTCAGGCACGTACAGGTCACGCAATTCT<br>GGAACGTTACGGTATGACCGAAACCAATATGAATACCAGCAATCCGTACGAAGGTA<br>AACGTATTGCAGGCACCGTTGGTTTTCCGCTGCCGGACGTTACCGTTCGTGTGACC<br>GATCCGGCAACAGGTCTGGCACTGCCTCCGGAACAGACAGGTATGATTGAAATTAA<br>AGGTCCGAACGTGTTCAAAGGCTATTGGCGTATGCCGGAAAAAACCGCAGCAGAAT<br>TTACCGCAGACGGCTTTTTTATCAGCGGTGATCTGGGTAAAATTGATCGTGACGGT<br>TACGTTCATATTGTTGGTCGTGGTAAAGACCTGGTGATTAGCGGTGGCTATAATAT<br>CTATCCGAAAGAAGTTGAAGGCGAGATCGATCAGATTGAAGGTGTTGTTGAAAGCG<br>CAGTTATTGGTGTTCCGCACCCGGATTTTGGTGAAGGTGTGACCGCAGTTGTTGTT<br>CGTAAACCGGGTGCAGCACTGGACGAAAAAGCAATTGTTAGCGCACTGCAAGATCG<br>CCTGGCACGTTATAAACAGCCGAAACGTATTATCTTTGCCGAAGATTTACCGCGTA<br>ATACGATGGGTAAAGTGCAGAAAAATATCCTGCGTCAGCAGTACGCCGATCTGTAT<br>ACACGTACCTAATAA |

**Table S2. List of primers used in this study.**

| Primer ID | Sequence (5' → 3') | Description | 5'-Modification |
|---|---|---|---|
| oDG127 | GGATTTGTCCTACTCAGGAG | Sequencing pSEVA insert FOR | |
| oDG128 | AAATCGTAATTATTGGGGAC | Sequencing pSEVA insert REV | |
| oDG172 | TTCATCTGATGCTGGATG | Sequencing downstream of KanR | |
| oDG176 | GCCTACTTCACCTATCCTGC | Sequencing upstream oriT | |
| oDG252 | ATCTTGGAACTACGCCTGCTAATACGAC | operon colony PCR FOR | |
| oDG253 | AACCGAGCGTTCTGAACAAATCCAG | operon colony PCR REV | |
| oDG254 | TGCGACACTGCATAACTGGACC | operon colony PCR from mNectarine REV | |
| oDG255 | CGCCAGGCATTTTCAGGTTTTTGG | operon colony PCR from mCardinal REV | |
| oDG256 | GGTTTTGCGCCATTCGATGGTGTC | Sequencing downstream of lacI promoter | |
| oDG304 | ACTAGTCTTGGACTCCTGTTGATAG | Amplification of pSEVA resistance FOR | Phosphorylation |
| oDG306 | TAGTAAATATAATCGTATAGAATAGGAGG | Amplification of fDG137 FOR | Phosphorylation |
| oDG307 | TTATTATTTGTCGTCGTCGTC | Amplification of fDG137 REV | Phosphorylation |
| oDG308 | CGAGACGATACAATAGAAGGAGG | Amplification of fDG138 FOR | Phosphorylation |
| oDG309 | TTATTACAGGTCTTCTTCGCTAATC | Amplification of fDG138 REV | Phosphorylation |
| oDG310 | AGATCGGACAAGGATCTAAGG | Amplification of fDG139 FOR | Phosphorylation |
| oDG311 | TTATTAGGTGCTATCCAGACC | Amplification of fDG139 REV | Phosphorylation |
| oDG312 | GAAAGAAAACAGCACTCCATAG | Amplification of fDG140 FOR | Phosphorylation |
| oDG313 | TTATTATGCATAATCCGGAAC | Amplification of fDG140 REV | Phosphorylation |
| oDG314 | ATTATAGGTCAAAAAATAGAGGAGG | Amplification of fDG141 FOR | Phosphorylation |
| oDG315 | TTATTATTTTTCAAACTGCGG | Amplification of fDG141 REV | Phosphorylation |
| oDG316 | AGGCAGTTCGCCAGTAAG | Amplification of fDG142 FOR | Phosphorylation |
| oDG317 | TTATTAACCCATTTGCTGTCC | Amplification of fDG142 REV | Phosphorylation |
| oDG318 | TATTTTAATAGGAGTCAGTAAGGAGAATAC | Amplification of fDG143 FOR | Phosphorylation |
| oDG319 | TTATTATTTACCCAGACGGTTC | Amplification of fDG143 REV | Phosphorylation |
| oDG382 | TTAATTAAAGGCATCAAATAAAACG | Amplification of pSEVA ori REV | Phosphorylation |
| oDG387 | ACTAGTCTTGGACTCCTGTTGA | Amplification of pSEVA backbone FOR | Phosphorylation |
| oDG411 | AGCGGATAACAATTTCACACAG | Amplification of pSEVA MCS FOR | Phosphorylation |
| oDG412 | CGCCAGGGTTTTCCC | Amplification of pSEVA MCS REV | Phosphorylation |
| oDG413 | ATTTTTTCGGTATATCCATCCT | Amplification of pSEVA ori FOR | Phosphorylation |
| oDG414 | CGCTATAATGACCCCGAA | Amplification of pSEVA resistance REV | Phosphorylation |
| oDG419 | GAAGGGCAATCAGCTGTTGC | Sequencing downstream lacI | |
| oDG424 | AGAAGATTGTGCACCGCC | Sequencing upstream lacI | |
| oDG425 | TAGGGTTTTCCCTCC | Sequencing upstream mCardinal | |
| oDG426 | CTCCCCCTCGCCTTC | Sequencing upstream mNectarine | |
| oDG427 | TTCGCCTTCTCCTTC | Sequencing upstream LSSmOrange | |
| oDG428 | TACCTGTACCCTGAC | Sequencing upstream mNeonGreen | |
| oDG429 | ATAGGTTGCATCACC | Sequencing upstream mT-Sapphire | |
| oDG430 | CGTGGCATCACCCTC | Sequencing upstream mTurquoise2 | |
| oDG431 | ACGGTTTACCTTCAC | Sequencing upstream mTagBFP2 | |
| oDG482 | CTCAAACCGTCACATTAGGG | Amplification of fDG200 FOR | Phosphorylation |
| oDG483 | GCTCCTAGCATATAAGGAGAAT | Amplification of fDG201 FOR | Phosphorylation |
| oDG484 | TAAATTCTTATCATTAAAATAAGGAGAG | Amplification of fDG202 FOR | Phosphorylation |
| oDG485 | TTATTAGTTTTTATACGGCAGGG | Amplification of fDG202 REV | Phosphorylation |
| oDG486 | GCTTAAACTCTCATAAGGAGGTA | Amplification of fDG203 FOR | Phosphorylation |
| oDG487 | TTATTAGGTGGCGTTACCAAC | Amplification of fDG203 REV | Phosphorylation |
| oDG488 | CTATCACTGCCCCGCTTTC | Amplification of fDG197 FOR | Phosphorylation |
| oDG492 | GCTGTGTGCACGAACC | Sequencing pSEVAx9x pBR322/rop ori REV | |
| oDG493 | CTAATCCTGTTACCAGTGGC | Sequencing pSEVAx9x pBR322/rop ori FOR | |
| oDG494 | TTTCTCCTTACGCATCTGTG | Sequencing pSEVAx9x pBR322/rop ori FOR | |
| oDG495 | ATTTCATCGGTCACATCGTT | Sequencing pSEVA2xx kanR REV | |
| oDG496 | GCGACCATTTATCGTCTGTA | Sequencing pSEVA2xx kanR FOR | |
| oDG498 | CACTCGCAATCAAATTCAGC | Sequencing upstream lacI | |
| oDG499 | CATGTGTCAGAGGTTTTCAC | Sequencing downstram rop | |
| oDG500 | TAACAGCGATCGTGTGTTTC | Sequencing downstram kanR | |
| oDG536 | ACTGAAATCACAGACTCAAC | Amplification of fDG210 FOR | Phosphorylation |
| oDG537 | TTATCACTGTTCGCTTTCG | Amplification of fDG210 REV | Phosphorylation |
| oDG538 | ACTTCGTCAATCGTACTCATAAG | Amplification of fDG211 FOR | Phosphorylation |
| oDG539 | ACTCCAGACATTTTTAAGGC | Amplification of fDG212 FOR | Phosphorylation |

| | | | |
|---|---|---|---|
| oDG540 | TTATTAGGCAATACCGTTCAG | Amplification of fDG212 REV | Phosphorylation |
| oDG541 | ACAAGACATTTAATTAAGGACAGG | Amplification of fDG213 FOR | Phosphorylation |
| oDG542 | TTATTAGATACGATCTTTCCAAACG | Amplification of fDG213 REV | Phosphorylation |
| oDG543 | GAATTTAAGCGCTTAAGGAGAG | Amplification of fDG214 FOR | Phosphorylation |
| oDG544 | TTATTAGGTACGTGTATACAGATCG | Amplification of fDG214 REV | Phosphorylation |
| oDG581 | CTGCACCGTTGGTAAC | Sequencing bauA middle REV | |
| oDG641 | GGTAAACAACTGGCGGTATGGATGC | Operon colony PCR FOR | |
| oDG642 | CAACCGAGCGTTCTGAACAAATCC | Operon colony PCR REV | |

**Table S3. List of LCR bridging oligos used in this study.**

| Bridging oligo ID | Bridge between: Upstream fragment ID | Bridge between: Downstream fragment ID | Sequence (5' → 3') |
|---|---|---|---|
| oDG320 | fDG197 | fDG137 | CTCACTATAGGGGAATTGTGAGCGGATAACAATTCCTAGTAAATATAATCGTATAGAATAGGAGGAACCATGGTCAGCA |
| oDG321 | fDG197 | fDG138 | CTCACTATAGGGGAATTGTGAGCGGATAACAATTCCCGAGACGATACAATAGAAGGAGGAGGCACACA |
| oDG322 | fDG197 | fDG139 | CTCACTATAGGGGAATTGTGAGCGGATAACAATTCCAGATCGGACAAGGATCTAAGGAAGAGAGGTTAAGAATG |
| oDG323 | fDG197 | fDG140 | CTCACTATAGGGGAATTGTGAGCGGATAACAATTCCGAAAGAAAACAGCACTCCATAGAAGGAGGTTTATATGGTC |
| oDG324 | fDG197 | fDG141 | CTCACTATAGGGGAATTGTGAGCGGATAACAATTCCATTATAGGTCAAAAAATAGAGGAGGACGGAAATCAATGGT |
| oDG325 | fDG197 | fDG142 | CTCACTATAGGGGAATTGTGAGCGGATAACAATTCCAGGCAGTTCGCCAGTAAGTAAGGAGGTAATC |
| oDG326 | fDG197 | fDG143 | CTCACTATAGGGGAATTGTGAGCGGATAACAATTCCTATTTTAATAGGAGTCAGTAAGGAGAATACGGATGGTCAGC |
| oDG327 | fDG137 | fDG138 | AGCACCGATTACAAAGACGACGACGACAAATAATAACGAGACGATACAATAGAAGGAGGAGGCACACA |
| oDG328 | fDG137 | fDG139 | AGCACCGATTACAAAGACGACGACGACAAATAATAAAGATCGGACAAGGATCTAAGGAAGAGAGGTTAAGAATG |
| oDG329 | fDG137 | fDG140 | AGCACCGATTACAAAGACGACGACGACAAATAATAAGAAAGAAAACAGCACTCCATAGAAGGAGGTTTATATGGTC |
| oDG330 | fDG137 | fDG141 | AGCACCGATTACAAAGACGACGACGACAAATAATAAATTATAGGTCAAAAAATAGAGGAGGACGGAAATCAATGGT |
| oDG331 | fDG137 | fDG142 | AGCACCGATTACAAAGACGACGACGACAAATAATAAAGGCAGTTCGCCAGTAAGTAAGGAGGTAATC |
| oDG332 | fDG137 | fDG143 | AGCACCGATTACAAAGACGACGACGACAAATAATAATATTTTAATAGGAGTCAGTAAGGAGAATACGGATGGTCAGC |
| oDG333 | fDG137 | fDG163 | AGCACCGATTACAAAGACGACGACGACAAATAATAAACTAGTCTTGGACTCCTGTTGATAGATCCAGTAATGAC |
| oDG334 | fDG138 | fDG139 | CAACTGAACAAAAACTGATTAGCGAAGAAGACCTGTAATAAAGATCGGACAAGGATCTAAGGAAGAGAGGTTAAGAATG |
| oDG335 | fDG138 | fDG140 | CAACTGAACAAAAACTGATTAGCGAAGAAGACCTGTAATAAGAAAGAAAACAGCACTCCATAGAAGGAGGTTTATATGGTC |
| oDG336 | fDG138 | fDG141 | CAACTGAACAAAAACTGATTAGCGAAGAAGACCTGTAATAAATTATAGGTCAAAAAATAGAGGAGGACGGAAATCAATGGT |
| oDG337 | fDG138 | fDG142 | CAACTGAACAAAAACTGATTAGCGAAGAAGACCTGTAATAAAGGCAGTTCGCCAGTAAGTAAGGAGGTAATC |
| oDG338 | fDG138 | fDG143 | CAACTGAACAAAAACTGATTAGCGAAGAAGACCTGTAATAATATTTTAATAGGAGTCAGTAAGGAGAATACGGATGGTCAGC |
| oDG339 | fDG138 | fDG163 | CAACTGAACAAAAACTGATTAGCGAAGAAGACCTGTAATAAACTAGTCTTGGACTCCTGTTGATAGATCCAGTAATGAC |
| oDG340 | fDG138 | fDG137 | CAACTGAACAAAAACTGATTAGCGAAGAAGACCTGTAATAATAGTAAATATAATCGTATAGAATAGGAGGAACCATGGTCAGCA |
| oDG341 | fDG139 | fDG140 | CGCTGCTGGGTCTGGATAGCACCTAATAAGAAAGAAAACAGCACTCCATAGAAGGAGGTTTATATGGTC |
| oDG342 | fDG139 | fDG141 | CGCTGCTGGGTCTGGATAGCACCTAATAAATTATAGGTCAAAAAATAGAGGAGGACGGAAATCAATGGT |
| oDG343 | fDG139 | fDG142 | CGCTGCTGGGTCTGGATAGCACCTAATAAAGGCAGTTCGCCAGTAAGTAAGGAGGTAATC |
| oDG344 | fDG139 | fDG143 | CGCTGCTGGGTCTGGATAGCACCTAATAATATTTTAATAGGAGTCAGTAAGGAGAATACGGATGGTCAGC |
| oDG345 | fDG139 | fDG163 | CGCTGCTGGGTCTGGATAGCACCTAATAAACTAGTCTTGGACTCCTGTTGATAGATCCAGTAATGAC |
| oDG346 | fDG139 | fDG137 | CGCTGCTGGGTCTGGATAGCACCTAATAATAGTAAATATAATCGTATAGAATAGGAGGAACCATGGTCAGCA |
| oDG347 | fDG139 | fDG138 | CGCTGCTGGGTCTGGATAGCACCTAATAACGAGACGATACAATAGAAGGAGGAGGCACACA |
| oDG348 | fDG140 | fDG141 | CGACCTATCCGTATGATGTTCCGGATTATGCATAATAAATTATAGGTCAAAAAATAGAGGAGGACGGAAATCAATGGT |
| oDG349 | fDG140 | fDG142 | CGACCTATCCGTATGATGTTCCGGATTATGCATAATAAAGGCAGTTCGCCAGTAAGTAAGGAGGTAATC |
| oDG350 | fDG140 | fDG143 | CGACCTATCCGTATGATGTTCCGGATTATGCATAATAATATTTTAATAGGAGTCAGTAAGGAGAATACGGATGGTCAGC |
| oDG351 | fDG140 | fDG163 | CGACCTATCCGTATGATGTTCCGGATTATGCATAATAAACTAGTCTTGGACTCCTGTTGATAGATCCAGTAATGAC |
| oDG352 | fDG140 | fDG137 | CGACCTATCCGTATGATGTTCCGGATTATGCATAATAATAGTAAATATAATCGTATAGAATAGGAGGAACCATGGTCAGCA |
| oDG353 | fDG140 | fDG138 | CGACCTATCCGTATGATGTTCCGGATTATGCATAATAACGAGACGATACAATAGAAGGAGGAGGCACACA |
| oDG354 | fDG140 | fDG139 | CGACCTATCCGTATGATGTTCCGGATTATGCATAATAAAGATCGGACAAGGATCTAAGGAAGAGAGGTTAAGAATG |
| oDG355 | fDG141 | fDG142 | ACAAATTGGAGCCATCCGCAGTTTGAAAAATAATAAAGGCAGTTCGCCAGTAAGTAAGGAGGTAATC |
| oDG356 | fDG141 | fDG143 | ACAAATTGGAGCCATCCGCAGTTTGAAAAATAATAATATTTTAATAGGAGTCAGTAAGGAGAATACGGATGGTCAGC |

| | | | |
|---|---|---|---|
| oDG357 | fDG141 | fDG163 | ACAAATTGGAGCCATCCGCAGTTTGAAAAATAATAAACTAGTCTTGGACTCCTGTTGATAGATCCAGTAATGAC |
| oDG358 | fDG141 | fDG137 | ACAAATTGGAGCCATCCGCAGTTTGAAAAATAATAATAGTAAATATAATCGTATAGAATAGGAGGAACCATGGTCAGCA |
| oDG359 | fDG141 | fDG138 | ACAAATTGGAGCCATCCGCAGTTTGAAAAATAATAACGAGACGATACAATAGAAGGAGGAGGCACACA |
| oDG360 | fDG141 | fDG139 | ACAAATTGGAGCCATCCGCAGTTTGAAAAATAATAAAGATCGGACAAGGATCTAAGGAAGAGAGGTTAAGAATG |
| oDG361 | fDG141 | fDG140 | ACAAATTGGAGCCATCCGCAGTTTGAAAAATAATAAGAAAGAAAACAGCACTCCATAGAAGGAGGTTTATATGGTC |
| oDG362 | fDG142 | fDG143 | GCATGACTGGTGGACAGCAAATGGGTTAATAATATTTTAATAGGAGTCAGTAAGGAGAATACGGATGGTCAGC |
| oDG363 | fDG142 | fDG163 | GCATGACTGGTGGACAGCAAATGGGTTAATAAACTAGTCTTGGACTCCTGTTGATAGATCCAGTAATGAC |
| oDG364 | fDG142 | fDG137 | GCATGACTGGTGGACAGCAAATGGGTTAATAATAGTAAATATAATCGTATAGAATAGGAGGAACCATGGTCAGCA |
| oDG365 | fDG142 | fDG138 | GCATGACTGGTGGACAGCAAATGGGTTAATAACGAGACGATACAATAGAAGGAGGAGGCACACA |
| oDG366 | fDG142 | fDG139 | GCATGACTGGTGGACAGCAAATGGGTTAATAAAGATCGGACAAGGATCTAAGGAAGAGAGGTTAAGAATG |
| oDG367 | fDG142 | fDG140 | GCATGACTGGTGGACAGCAAATGGGTTAATAAGAAAGAAAACAGCACTCCATAGAAGGAGGTTTATATGGTC |
| oDG368 | fDG142 | fDG141 | GCATGACTGGTGGACAGCAAATGGGTTAATAAATTATAGGTCAAAAAATAGAGGAGGACGGAAATCAATGGT |
| oDG369 | fDG143 | fDG163 | GTACCTATACCGATATTGAAATGAACCGTCTGGGTAAATAATAAACTAGTCTTGGACTCCTGTTGATAGATCCAGTAATGAC |
| oDG370 | fDG143 | fDG137 | GTACCTATACCGATATTGAAATGAACCGTCTGGGTAAATAATAATAGTAAATATAATCGTATAGAATAGGAGGAACCATGGTCAGCA |
| oDG371 | fDG143 | fDG138 | GTACCTATACCGATATTGAAATGAACCGTCTGGGTAAATAATAACGAGACGATACAATAGAAGGAGGAGGCACACA |
| oDG372 | fDG143 | fDG139 | GTACCTATACCGATATTGAAATGAACCGTCTGGGTAAATAATAAAGATCGGACAAGGATCTAAGGAAGAGAGGTTAAGAATG |
| oDG373 | fDG143 | fDG140 | GTACCTATACCGATATTGAAATGAACCGTCTGGGTAAATAATAAGAAAGAAAACAGCACTCCATAGAAGGAGGTTTATATGGTC |
| oDG374 | fDG143 | fDG141 | GTACCTATACCGATATTGAAATGAACCGTCTGGGTAAATAATAAATTATAGGTCAAAAAATAGAGGAGGACGGAAATCAATGGT |
| oDG375 | fDG143 | fDG142 | GTACCTATACCGATATTGAAATGAACCGTCTGGGTAAATAATAAAGGCAGTTCGCCAGTAAGTAAGGAGGTAATC |
| oDG417 | fDG163 | fDG164 | GCATAACCCTGCTTCGGGGTCATTATAGCGATTTTTTCGGTATATCCATCCTTTTTCGCACGATATACAG |
| oDG474 | fDG164 | fDG201 | TCGACTGAGCCTTTCGTTTTATTTGATGCCTTTAATTAAGCTCCTAGCATATAAGGAGAATTTTTATGAACAAAGGTGTTATG |
| oDG475 | fDG197 | fDG201 | CTCACTATAGGGGAATTGTGAGCGGATAACAATTCCGCTCCTAGCATATAAGGAGAATTTTTATGAACAAAGGTGTTATG |
| oDG476 | fDG164 | fDG202 | TCGACTGAGCCTTTCGTTTTATTTGATGCCTTTAATTAATAAATTCTTATCATTAAAATAAGGAGAGTCTCTATGCATCATCACC |
| oDG477 | fDG197 | fDG202 | CTCACTATAGGGGAATTGTGAGCGGATAACAATTCCTAAATTCTTATCATTAAAATAAGGAGAGTCTCTATGCATCATCACC |
| oDG478 | fDG202 | fDG163 | CTATATCCTGAAAGAAATTGATACCCTGCCGTATAAAAACTAATAAACTAGTCTTGGACTCCTGTTGATAGATCCAGTAATGAC |
| oDG489 | fDG164 | fDG197 | TCGACTGAGCCTTTCGTTTTATTTGATGCCTTTAATTAACTATCACTGCCCGCTTTCCAGTCGGG |
| oDG519 | fDG201 | fDG202 | ACAAATTGGAGCCATCCGCAGTTTGAAAAATAATAATAAATTCTTATCATTAAAATAAGGAGAGTCTCTATGCATCATCACC |
| oDG520 | fDG202 | fDG137 | CTATATCCTGAAAGAAATTGATACCCTGCCGTATAAAAACTAATAATAGTAAATATAATCGTATAGAATAGGAGGAACCATGGTCAGCA |
| oDG521 | fDG202 | fDG138 | CTATATCCTGAAAGAAATTGATACCCTGCCGTATAAAAACTAATAACGAGACGATACAATAGAAGGAGGAGGCACACA |
| oDG522 | fDG202 | fDG140 | CTATATCCTGAAAGAAATTGATACCCTGCCGTATAAAAACTAATAAGAAAGAAAACAGCACTCCATAGAAGGAGGTTTATATGGTC |
| oDG523 | fDG202 | fDG142 | CTATATCCTGAAAGAAATTGATACCCTGCCGTATAAAAACTAATAAAGGCAGTTCGCCAGTAAGTAAGGAGGTAATC |
| oDG524 | fDG202 | fDG143 | CTATATCCTGAAAGAAATTGATACCCTGCCGTATAAAAACTAATAATATTTTAATAGGAGTCAGTAAGGAGAATACGGATGGTCAGC |
| oDG525 | fDG202 | fDG201 | CTATATCCTGAAAGAAATTGATACCCTGCCGTATAAAAACTAATAAGCTCCTAGCATATAAGGAGAATTTTTATGAACAAAGGTGTTATG |
| oDG526 | fDG137 | fDG202 | AGCACCGATTACAAAGACGACGACGACAAATAATAATAAATTCTTATCATTAAAATAAGGAGAGTCTCTATGCATCATCACC |
| oDG527 | fDG137 | fDG201 | AGCACCGATTACAAAGACGACGACGACAAATAATAAGCTCCTAGCATATAAGGAGAATTTTTATGAACAAAGGTGTTATG |
| oDG528 | fDG138 | fDG202 | CAACTGAACAAAAACTGATTAGCGAAGAAGACCTGTAATAATAAATTCTTATCATTAAAATAAGGAGAGTCTCTATGCATCATCACC |
| oDG529 | fDG138 | fDG201 | CAACTGAACAAAAACTGATTAGCGAAGAAGACCTGTAATAAGCTCCTAGCATATAAGGAGAATTTTTATGAACAAAGGTGTTATG |
| oDG530 | fDG140 | fDG202 | CGACCTATCCGTATGATGTTCCGGATTATGCATAATAATAAATTCTTATCATTAAAATAAGGAGAGTCTCTATGCATCATCACC |

| | | | |
|---|---|---|---|
| oDG531 | fDG140 | fDG201 | CGACCTATCCGTATGATGTTCCGGATTATGCATAATAAGCTCCTAGCATATAAGGAGAATTTTTATGAACAAAGGTGTTATG |
| oDG532 | fDG142 | fDG202 | GCATGACTGGTGGACAGCAAATGGGTTAATAATAAATTCTTATCATTAAAATAAGGAGAGTCTCTATGCATCATCACC |
| oDG533 | fDG142 | fDG201 | GCATGACTGGTGGACAGCAAATGGGTTAATAAGCTCCTAGCATATAAGGAGAATTTTTATGAACAAAGGTGTTATG |
| oDG534 | fDG143 | fDG202 | GTACCTATACCGATATTGAAATGAACCGTCTGGGTAAATAATAATAAATTCTTATCATTAAAATAAGGAGAGTCTCTATGCATCATCACC |
| oDG535 | fDG143 | fDG201 | GTACCTATACCGATATTGAAATGAACCGTCTGGGTAAATAATAAGCTCCTAGCATATAAGGAGAATTTTTATGAACAAAGGTGTTATG |
| oDG545 | fDG164 | fDG210 | TCGACTGAGCCTTTCGTTTTATTTGATGCCTTTAATTAAACTGAAATCACAGACTCAACTCATTTAAGCGCAG |
| oDG546 | fDG164 | fDG211 | TCGACTGAGCCTTTCGTTTTATTTGATGCCTTTAATTAAACTTCGTCAATCGTACTCATAAGGAGGTATATATGCTGC |
| oDG547 | fDG164 | fDG212 | TCGACTGAGCCTTTCGTTTTATTTGATGCCTTTAATTAAACTCCAGACATTTTTAAGGCAGGTTTTTATGCATCATC |
| oDG548 | fDG164 | fDG213 | TCGACTGAGCCTTTCGTTTTATTTGATGCCTTTAATTAAACAAGACATTTAATTAAGGACAGGTACGTTTATGCATCATC |
| oDG549 | fDG164 | fDG214 | TCGACTGAGCCTTTCGTTTTATTTGATGCCTTTAATTAAGAATTTAAGCGCTTAAGGAGAGTCATATATGCATCATCAC |
| oDG550 | fDG197 | fDG210 | CTCACTATAGGGGAATTGTGAGCGGATAACAATTCCACTGAAATCACAGACTCAACTCATTTAAGCGCAG |
| oDG551 | fDG197 | fDG211 | CTCACTATAGGGGAATTGTGAGCGGATAACAATTCCACTTCGTCAATCGTACTCATAAGGAGGTATATATGCTGC |
| oDG552 | fDG197 | fDG212 | CTCACTATAGGGGAATTGTGAGCGGATAACAATTCCACTCCAGACATTTTTAAGGCAGGTTTTTATGCATCATC |
| oDG553 | fDG197 | fDG213 | CTCACTATAGGGGAATTGTGAGCGGATAACAATTCCACAAGACATTTAATTAAGGACAGGTACGTTTATGCATCATC |
| oDG554 | fDG197 | fDG214 | CTCACTATAGGGGAATTGTGAGCGGATAACAATTCCGAATTTAAGCGCTTAAGGAGAGTCATATATGCATCATCAC |
| oDG555 | fDG210 | fDG163 | CAAAACGTTATACCGACGAAAGCGAACAGTGATAAACTAGTCTTGGACTCCTGTTGATAGATCCAGTAATGAC |
| oDG556 | fDG210 | fDG211 | CAAAACGTTATACCGACGAAAGCGAACAGTGATAAACTTCGTCAATCGTACTCATAAGGAGGTATATATGCTGC |
| oDG557 | fDG210 | fDG212 | CAAAACGTTATACCGACGAAAGCGAACAGTGATAAACTCCAGACATTTTTAAGGCAGGTTTTTATGCATCATC |
| oDG558 | fDG210 | fDG213 | CAAAACGTTATACCGACGAAAGCGAACAGTGATAAACAAGACATTTAATTAAGGACAGGTACGTTTATGCATCATC |
| oDG559 | fDG210 | fDG214 | CAAAACGTTATACCGACGAAAGCGAACAGTGATAAGAATTTAAGCGCTTAAGGAGAGTCATATATGCATCATCAC |
| oDG560 | fDG211 | fDG210 | CAACTGAACAAAAACTGATTAGCGAAGAAGACCTGTAATAAACTGAAATCACAGACTCAACTCATTTAAGCGCAG |
| oDG561 | fDG211 | fDG212 | CAACTGAACAAAAACTGATTAGCGAAGAAGACCTGTAATAAACTCCAGACATTTTTAAGGCAGGTTTTTATGCATCATC |
| oDG562 | fDG211 | fDG213 | CAACTGAACAAAAACTGATTAGCGAAGAAGACCTGTAATAAACAAGACATTTAATTAAGGACAGGTACGTTTATGCATCATC |
| oDG563 | fDG211 | fDG214 | CAACTGAACAAAAACTGATTAGCGAAGAAGACCTGTAATAAGAATTTAAGCGCTTAAGGAGAGTCATATATGCATCATCAC |
| oDG564 | fDG212 | fDG163 | GTTGGTGAAGCACTGAACGGTATTGCCTAATAAACTAGTCTTGGACTCCTGTTGATAGATCCAGTAATGAC |
| oDG565 | fDG212 | fDG210 | GTTGGTGAAGCACTGAACGGTATTGCCTAATAAACTGAAATCACAGACTCAACTCATTTAAGCGCAG |
| oDG566 | fDG212 | fDG211 | GTTGGTGAAGCACTGAACGGTATTGCCTAATAAACTTCGTCAATCGTACTCATAAGGAGGTATATATGCTGC |
| oDG567 | fDG212 | fDG213 | GTTGGTGAAGCACTGAACGGTATTGCCTAATAAACAAGACATTTAATTAAGGACAGGTACGTTTATGCATCATC |
| oDG568 | fDG212 | fDG214 | GTTGGTGAAGCACTGAACGGTATTGCCTAATAAGAATTTAAGCGCTTAAGGAGAGTCATATATGCATCATCAC |
| oDG569 | fDG213 | fDG163 | CGAATTTTGTAATATTCAGACCGTTTGGAAAGATCGTATCTAATAAACTAGTCTTGGACTCCTGTTGATAGATCCAGTAATGAC |
| oDG570 | fDG213 | fDG210 | CGAATTTTGTAATATTCAGACCGTTTGGAAAGATCGTATCTAATAAACTGAAATCACAGACTCAACTCATTTAAGCGCAG |
| oDG571 | fDG213 | fDG211 | CGAATTTTGTAATATTCAGACCGTTTGGAAAGATCGTATCTAATAAACTTCGTCAATCGTACTCATAAGGAGGTATATATGCTGC |
| oDG572 | fDG213 | fDG212 | CGAATTTTGTAATATTCAGACCGTTTGGAAAGATCGTATCTAATAAACTCCAGACATTTTTAAGGCAGGTTTTTATGCATCATC |
| oDG573 | fDG213 | fDG214 | CGAATTTTGTAATATTCAGACCGTTTGGAAAGATCGTATCTAATAAGAATTTAAGCGCTTAAGGAGAGTCATATATGCATCATCAC |
| oDG574 | fDG214 | fDG163 | CAGCAGTACGCCGATCTGTATACACGTACCTAATAAACTAGTCTTGGACTCCTGTTGATAGATCCAGTAATGAC |
| oDG575 | fDG214 | fDG210 | CAGCAGTACGCCGATCTGTATACACGTACCTAATAAACTGAAATCACAGACTCAACTCATTTAAGCGCAG |
| oDG576 | fDG214 | fDG211 | CAGCAGTACGCCGATCTGTATACACGTACCTAATAAACTTCGTCAATCGTACTCATAAGGAGGTATATATGCTGC |

| | | | |
|---|---|---|---|
| oDG577 | fDG214 | fDG212 | CAGCAGTACGCCGATCTGTATACACGTACCTAATAAACTCCAGACATTTTTAAGGCAGGTTTTTATGCATCATC |
| oDG578 | fDG214 | fDG213 | CAGCAGTACGCCGATCTGTATACACGTACCTAATAAACAAGACATTTAATTAAGGACAGGTACGTTTATGCATCATC |
| oDG605 | fDG210 | fDG138 | CAAAACGTTATACCGACGAAAGCGAACAGTGATAACGAGACGATACAATAGAAGGAGGAGGCACACA |
| oDG606 | fDG210 | fDG140 | CAAAACGTTATACCGACGAAAGCGAACAGTGATAAGAAAGAAAACAGCACTCCATAGAAGGAGGTTTATATGGTC |
| oDG607 | fDG210 | fDG142 | CAAAACGTTATACCGACGAAAGCGAACAGTGATAAAGGCAGTTCGCCAGTAAGTAAGGAGGTAATC |
| oDG608 | fDG210 | fDG143 | CAAAACGTTATACCGACGAAAGCGAACAGTGATAATATTTTAATAGGAGTCAGTAAGGAGAATACGGATGGTCAGC |
| oDG609 | fDG140 | fDG210 | CGACCTATCCGTATGATGTTCCGGATTATGCATAATAAACTGAAATCACAGACTCAACTCATTTAAGCGCAG |
| oDG610 | fDG142 | fDG210 | GCATGACTGGTGGACAGCAAATGGGTTAATAAACTGAAATCACAGACTCAACTCATTTAAGCGCAG |
| oDG611 | fDG143 | fDG210 | GTACCTATACCGATATTGAAATGAACCGTCTGGGTAAATAATAAACTGAAATCACAGACTCAACTCATTTAAGCGCAG |
| oDG612 | fDG137 | fDG211 | AGCACCGATTACAAAGACGACGACGACAAATAATAAACTTCGTCAATCGTACTCATAAGGAGGTATATATGCTGC |
| oDG613 | fDG140 | fDG211 | CGACCTATCCGTATGATGTTCCGGATTATGCATAATAAACTTCGTCAATCGTACTCATAAGGAGGTATATATGCTGC |
| oDG614 | fDG142 | fDG211 | GCATGACTGGTGGACAGCAAATGGGTTAATAAACTTCGTCAATCGTACTCATAAGGAGGTATATATGCTGC |
| oDG615 | fDG143 | fDG211 | GTACCTATACCGATATTGAAATGAACCGTCTGGGTAAATAATAAACTTCGTCAATCGTACTCATAAGGAGGTATATATGCTGC |
| oDG616 | fDG212 | fDG137 | GTTGGTGAAGCACTGAACGGTATTGCCTAATAATAGTAAATATAATCGTATAGAATAGGAGGAACCATGGTCAGCA |
| oDG617 | fDG212 | fDG138 | GTTGGTGAAGCACTGAACGGTATTGCCTAATAACGAGACGATACAATAGAAGGAGGAGGCACACA |
| oDG618 | fDG212 | fDG140 | GTTGGTGAAGCACTGAACGGTATTGCCTAATAAGAAAGAAAACAGCACTCCATAGAAGGAGGTTTATATGGTC |
| oDG619 | fDG212 | fDG142 | GTTGGTGAAGCACTGAACGGTATTGCCTAATAAAGGCAGTTCGCCAGTAAGTAAGGAGGTAATC |
| oDG620 | fDG212 | fDG143 | GTTGGTGAAGCACTGAACGGTATTGCCTAATAATATTTTAATAGGAGTCAGTAAGGAGAATACGGATGGTCAGC |
| oDG621 | fDG137 | fDG212 | AGCACCGATTACAAAGACGACGACGACAAATAATAAACTCCAGACATTTTTAAGGCAGGTTTTTATGCATCATC |
| oDG622 | fDG140 | fDG212 | CGACCTATCCGTATGATGTTCCGGATTATGCATAATAAACTCCAGACATTTTTAAGGCAGGTTTTTATGCATCATC |
| oDG623 | fDG142 | fDG212 | GCATGACTGGTGGACAGCAAATGGGTTAATAAACTCCAGACATTTTTAAGGCAGGTTTTTATGCATCATC |
| oDG624 | fDG143 | fDG212 | GTACCTATACCGATATTGAAATGAACCGTCTGGGTAAATAATAAACTCCAGACATTTTTAAGGCAGGTTTTTATGCATCATC |
| oDG625 | fDG213 | fDG137 | CGAATTTTGTAATATTCAGACCGTTTGGAAAGATCGTATCTAATAATAGTAAATATAATCGTATAGAATAGGAGGAACCATGGTCAGCA |
| oDG626 | fDG213 | fDG138 | CGAATTTTGTAATATTCAGACCGTTTGGAAAGATCGTATCTAATAACGAGACGATACAATAGAAGGAGGAGGCACACA |
| oDG627 | fDG213 | fDG142 | CGAATTTTGTAATATTCAGACCGTTTGGAAAGATCGTATCTAATAAAGGCAGTTCGCCAGTAAGTAAGGAGGTAATC |
| oDG628 | fDG213 | fDG143 | CGAATTTTGTAATATTCAGACCGTTTGGAAAGATCGTATCTAATAATATTTTAATAGGAGTCAGTAAGGAGAATACGGATGGTCAGC |
| oDG629 | fDG137 | fDG213 | AGCACCGATTACAAAGACGACGACGACAAATAATAAACAAGACATTTAATTAAGGACAGGTACGTTTATGCATCATC |
| oDG630 | fDG142 | fDG213 | GCATGACTGGTGGACAGCAAATGGGTTAATAAACAAGACATTTAATTAAGGACAGGTACGTTTATGCATCATC |
| oDG631 | fDG143 | fDG213 | GTACCTATACCGATATTGAAATGAACCGTCTGGGTAAATAATAAACAAGACATTTAATTAAGGACAGGTACGTTTATGCATCATC |
| oDG632 | fDG214 | fDG137 | CAGCAGTACGCCGATCTGTATACACGTACCTAATAATAGTAAATATAATCGTATAGAATAGGAGGAACCATGGTCAGCA |
| oDG633 | fDG214 | fDG138 | CAGCAGTACGCCGATCTGTATACACGTACCTAATAACGAGACGATACAATAGAAGGAGGAGGCACACA |
| oDG634 | fDG214 | fDG140 | CAGCAGTACGCCGATCTGTATACACGTACCTAATAAGAAAGAAAACAGCACTCCATAGAAGGAGGTTTATATGGTC |
| oDG635 | fDG214 | fDG143 | CAGCAGTACGCCGATCTGTATACACGTACCTAATAATATTTTAATAGGAGTCAGTAAGGAGAATACGGATGGTCAGC |
| oDG636 | fDG137 | fDG214 | AGCACCGATTACAAAGACGACGACGACAAATAATAAGAATTTAAGCGCTTAAGGAGAGTCATATATGCATCATCAC |
| oDG637 | fDG140 | fDG214 | CGACCTATCCGTATGATGTTCCGGATTATGCATAATAAGAATTTAAGCGCTTAAGGAGAGTCATATATGCATCATCAC |
| oDG638 | fDG143 | fDG214 | GTACCTATACCGATATTGAAATGAACCGTCTGGGTAAATAATAAGAATTTAAGCGCTTAAGGAGAGTCATATATGCATCATCAC |

**Table S4. List of plasmids used in this study.** The operon designs corresponding to each plasmid are abbreviated as string of letters representing the order of assembled genes (C: mCardinal, N: mNectarine, O: LSSmOrange, G: mNeonGreen, S: mT-Sapphire, T: mTurquoise2, B: mTagBFP2, X: XylE, D: DocS, A: AspA, P: PanD, U: BauA, –: 'empty vector'). Plasmid sequences result from the concatenation of the fragment sequences in the listed fragment composition order (see **Table S1**).

| **Plasmid ID** | **Operon design** | **Description** | **Fragment composition** |
|---|---|---|---|
| pSEVA291 | – | Template for SEVA plasmid backbone[37] | – |
| pDG072 | – | pSEVA29x_lacI_pT7lacO_mcs | fDG197, fDG199, fDG164 |
| vDG-L-001 | C | pSEVA29x_lacI_pT7lacO_C | fDG197, fDG137, fDG163, fDG164 |
| vDG-L-002 | N | pSEVA29x_lacI_pT7lacO_N | fDG197, fDG138, fDG163, fDG164 |
| vDG-L-003 | O | pSEVA29x_lacI_pT7lacO_O | fDG197, fDG139, fDG163, fDG164 |
| vDG-L-004 | G | pSEVA29x_lacI_pT7lacO_G | fDG197, fDG140, fDG163, fDG164 |
| vDG-L-005 | S | pSEVA29x_lacI_pT7lacO_S | fDG197, fDG141, fDG163, fDG164 |
| vDG-L-006 | T | pSEVA29x_lacI_pT7lacO_T | fDG197, fDG142, fDG163, fDG164 |
| vDG-L-007 | B | pSEVA29x_lacI_pT7lacO_B | fDG197, fDG143, fDG163, fDG164 |
| pDGogo3o7-018 | CSG | pSEVA29x_lacI_pT7lacO_CSG | fDG197, fDG137, fDG141, fDG140, fDG163, fDG164 |
| pDGogo3o7-020 | CSB | pSEVA29x_lacI_pT7lacO_CSB | fDG197, fDG137, fDG141, fDG143, fDG163, fDG164 |
| pDGogo3o7-022 | CTO | pSEVA29x_lacI_pT7lacO_CTO | fDG197, fDG137, fDG142, fDG139, fDG163, fDG164 |
| pDGogo3o7-042 | NGO | pSEVA29x_lacI_pT7lacO_NGO | fDG197, fDG138, fDG140, fDG139, fDG163, fDG164 |
| pDGogo3o7-054 | NTS | pSEVA29x_lacI_pT7lacO_NTS | fDG197, fDG138, fDG142, fDG141, fDG163, fDG164 |
| pDGogo3o7-055 | NTB | pSEVA29x_lacI_pT7lacO_NTB | fDG197, fDG138, fDG142, fDG143, fDG163, fDG164 |
| pDGogo3o7-056 | NBC | pSEVA29x_lacI_pT7lacO_NBC | fDG197, fDG138, fDG143, fDG137, fDG163, fDG164 |
| pDGogo3o7-057 | NBO | pSEVA29x_lacI_pT7lacO_NBO | fDG197, fDG138, fDG143, fDG139, fDG163, fDG164 |
| pDGogo3o7-064 | OCT | pSEVA29x_lacI_pT7lacO_OCT | fDG197, fDG139, fDG137, fDG142, fDG163, fDG164 |
| pDGogo3o7-077 | OSN | pSEVA29x_lacI_pT7lacO_OSN | fDG197, fDG139, fDG141, fDG138, fDG163, fDG164 |
| pDGogo3o7-094 | GCT | pSEVA29x_lacI_pT7lacO_GCT | fDG197, fDG140, fDG137, fDG142, fDG163, fDG164 |
| pDGogo3o7-100 | GNB | pSEVA29x_lacI_pT7lacO_GNB | fDG197, fDG140, fDG138, fDG143, fDG163, fDG164 |
| pDGogo3o7-101 | GOC | pSEVA29x_lacI_pT7lacO_GOC | fDG197, fDG140, fDG139, fDG137, fDG163, fDG164 |
| pDGogo3o7-102 | GON | pSEVA29x_lacI_pT7lacO_GON | fDG197, fDG140, fDG139, fDG138, fDG163, fDG164 |
| pDGogo3o7-114 | GTS | pSEVA29x_lacI_pT7lacO_GTS | fDG197, fDG140, fDG142, fDG141, fDG163, fDG164 |
| vDG-L-022 | GTB | pSEVA29x_lacI_pT7lacO_GTB | fDG197, fDG140, fDG142, fDG143, fDG163, fDG164 |
| pDGogo3o7-122 | SCO | pSEVA29x_lacI_pT7lacO_SCO | fDG197, fDG141, fDG137, fDG139, fDG163, fDG164 |
| pDGogo3o7-128 | SNG | pSEVA29x_lacI_pT7lacO_SNG | fDG197, fDG141, fDG138, fDG140, fDG163, fDG164 |
| pDGogo3o7-129 | SNT | pSEVA29x_lacI_pT7lacO_SNT | fDG197, fDG141, fDG138, fDG142, fDG163, fDG164 |
| pDGogo3o7-136 | SGC | pSEVA29x_lacI_pT7lacO_SGC | fDG197, fDG141, fDG140, fDG137, fDG163, fDG164 |
| pDGogo3o7-139 | SGT | pSEVA29x_lacI_pT7lacO_SGT | fDG197, fDG141, fDG140, fDG142, fDG163, fDG164 |
| pDGogo3o7-144 | STG | pSEVA29x_lacI_pT7lacO_STG | fDG197, fDG141, fDG142, fDG140, fDG163, fDG164 |
| pDGogo3o7-154 | TCS | pSEVA29x_lacI_pT7lacO_TCS | fDG197, fDG142, fDG137, fDG141, fDG163, fDG164 |
| pDGogo3o7-155 | TCB | pSEVA29x_lacI_pT7lacO_TCB | fDG197, fDG142, fDG137, fDG143, fDG163, fDG164 |
| vDG-L-026 | TBC | pSEVA29x_lacI_pT7lacO_TBC | fDG197, fDG142, fDG143, fDG137, fDG163, fDG164 |
| pDGogo3o7-177 | TBN | pSEVA29x_lacI_pT7lacO_TBN | fDG197, fDG142, fDG143, fDG138, fDG163, fDG164 |
| pDGogo3o7-182 | BCO | pSEVA29x_lacI_pT7lacO_BCO | fDG197, fDG143, fDG137, fDG139, fDG163, fDG164 |
| pDGogo3o7-185 | BCT | pSEVA29x_lacI_pT7lacO_BCT | fDG197, fDG143, fDG137, fDG142, fDG163, fDG164 |
| pDGogo3o7-193 | BOG | pSEVA29x_lacI_pT7lacO_BOG | fDG197, fDG143, fDG139, fDG140, fDG163, fDG164 |
| pDGogo3o7-200 | BGT | pSEVA29x_lacI_pT7lacO_BGT | fDG197, fDG143, fDG140, fDG142, fDG163, fDG164 |
| pDGogo3o7-206 | BTC | pSEVA29x_lacI_pT7lacO_BTC | fDG197, fDG143, fDG142, fDG137, fDG163, fDG164 |
| pDGogo3o7-210 | BTS | pSEVA29x_lacI_pT7lacO_BTS | fDG197, fDG143, fDG142, fDG141, fDG163, fDG164 |
| vDG-L-013 | CNOGSTB | pSEVA29x_lacI_pT7lacO_CNOGSTB | fDG197, fDG137, fDG138, fDG139, fDG140, fDG141, fDG142, fDG143, fDG163, fDG164 |
| vDG-L-012 | CNOGST | pSEVA29x_lacI_pT7lacO_CNOGST | fDG197, fDG137, fDG138, fDG139, fDG140, fDG141, fDG142, fDG163, fDG164 |
| vDG-L-011 | CNOGS | pSEVA29x_lacI_pT7lacO_CNOGS | fDG197, fDG137, fDG138, fDG139, fDG140, fDG141, fDG163, fDG164 |
| vDG-L-010 | CNOG | pSEVA29x_lacI_pT7lacO_CNOG | fDG197, fDG137, fDG138, fDG139, fDG140, fDG163, fDG164 |
| vDG-L-009 | CNO | pSEVA29x_lacI_pT7lacO_CNO | fDG197, fDG137, fDG138, fDG139, fDG163, fDG164 |
| vDG-L-008 | CN | pSEVA29x_lacI_pT7lacO_CN | fDG197, fDG137, fDG138, fDG163, fDG164 |
| vDG-L-014 | CNG | pSEVA29x_lacI_pT7lacO_CNG | fDG197, fDG137, fDG138, fDG140, fDG163, fDG164 |
| vDG-L-015 | CNGT | pSEVA29x_lacI_pT7lacO_CNGT | fDG197, fDG137, fDG138, fDG140, fDG142, fDG163, fDG164 |
| vDG-L-016 | CNGTB | pSEVA29x_lacI_pT7lacO_CNGTB | fDG197, fDG137, fDG138, fDG140, fDG142, fDG143, fDG163, fDG164 |
| vDG-L-017 | NG | pSEVA29x_lacI_pT7lacO_NG | fDG197, fDG138, fDG140, fDG163, fDG164 |
| vDG-L-018 | NGT | pSEVA29x_lacI_pT7lacO_NGT | fDG197, fDG138, fDG140, fDG142, fDG163, fDG164 |
| vDG-L-019 | NGTB | pSEVA29x_lacI_pT7lacO_NGTB | fDG197, fDG138, fDG140, fDG142, fDG143, fDG163, fDG164 |
| vDG-L-020 | NGTBC | pSEVA29x_lacI_pT7lacO_NGTBC | fDG197, fDG138, fDG140, fDG142, fDG143, fDG137, fDG163, fDG164 |
| vDG-L-021 | GT | pSEVA29x_lacI_pT7lacO_GT | fDG197, fDG140, fDG142, fDG163, fDG164 |
| vDG-L-023 | GTBC | pSEVA29x_lacI_pT7lacO_GTBC | fDG197, fDG140, fDG142, fDG143, fDG137, fDG163, fDG164 |
| vDG-L-024 | GTBCN | pSEVA29x_lacI_pT7lacO_GTBCN | fDG197, fDG140, fDG142, fDG143, fDG137, fDG138, fDG163, fDG164 |
| vDG-L-025 | TB | pSEVA29x_lacI_pT7lacO_TB | fDG197, fDG142, fDG143, fDG163, fDG164 |

| | | | |
|---|---|---|---|
| vDG-L-027 | TBCN | pSEVA29x_lacI_pT7lacO_TBCN | fDG197, fDG142, fDG143, fDG137, fDG138, fDG163, fDG164 |
| vDG-L-028 | TBCNG | pSEVA29x_lacI_pT7lacO_TBCNG | fDG197, fDG142, fDG143, fDG137, fDG138, fDG140, fDG163, fDG164 |
| vDG-L-029 | BC | pSEVA29x_lacI_pT7lacO_BC | fDG197, fDG143, fDG137, fDG163, fDG164 |
| vDG-L-030 | BCN | pSEVA29x_lacI_pT7lacO_BCN | fDG197, fDG143, fDG137, fDG138, fDG163, fDG164 |
| vDG-L-031 | BCNG | pSEVA29x_lacI_pT7lacO_BCNG | fDG197, fDG143, fDG137, fDG138, fDG140, fDG163, fDG164 |
| vDG-L-032 | BCNGT | pSEVA29x_lacI_pT7lacO_BCNGT | fDG197, fDG143, fDG137, fDG138, fDG140, fDG142, fDG163, fDG164 |
| v4FP-rand-020 | CTNB | pSEVA29x_lacI_pT7lacO_CTNB | fDG197, fDG137, fDG142, fDG138, fDG143, fDG163, fDG164 |
| v4FP-rand-068 | GTCB | pSEVA29x_lacI_pT7lacO_GTCB | fDG197, fDG140, fDG142, fDG137, fDG143, fDG163, fDG164 |
| v4FP-025-064 | GBNT | pSEVA29x_lacI_pT7lacO_GBNT | fDG197, fDG140, fDG143, fDG138, fDG142, fDG163, fDG164 |
| v4FP-seq02-1-029 | NCTG | pSEVA29x_lacI_pT7lacO_NCTG | fDG197, fDG138, fDG137, fDG142, fDG140, fDG163, fDG164 |
| v4FP-rand-024 | CTBG | pSEVA29x_lacI_pT7lacO_CTBG | fDG197, fDG137, fDG142, fDG143, fDG140, fDG163, fDG164 |
| v4FP-rand-074 | BCNT | pSEVA29x_lacI_pT7lacO_BCNT | fDG197, fDG143, fDG137, fDG138, fDG142, fDG163, fDG164 |
| v4FP-rand-088 | BGNT | pSEVA29x_lacI_pT7lacO_BGNT | fDG197, fDG143, fDG140, fDG138, fDG142, fDG163, fDG164 |
| v4FP-seq02-1-032 | NGCT | pSEVA29x_lacI_pT7lacO_NGCT | fDG197, fDG138, fDG140, fDG137, fDG142, fDG163, fDG164 |
| v4FP-rand-078 | BCTG | pSEVA29x_lacI_pT7lacO_BCTG | fDG197, fDG143, fDG137, fDG142, fDG140, fDG163, fDG164 |
| v4FP-025-102 | TCBG | pSEVA29x_lacI_pT7lacO_TCBG | fDG197, fDG142, fDG137, fDG143, fDG140, fDG163, fDG164 |
| v4FP-rand-041 | NBTC | pSEVA29x_lacI_pT7lacO_NBTC | fDG197, fDG138, fDG143, fDG142, fDG137, fDG163, fDG164 |
| v4FP-rand-079 | BNCG | pSEVA29x_lacI_pT7lacO_BNCG | fDG197, fDG143, fDG138, fDG137, fDG140, fDG163, fDG164 |
| v4FP-025-113 | TGBC | pSEVA29x_lacI_pT7lacO_TGBC | fDG197, fDG142, fDG140, fDG143, fDG137, fDG163, fDG164 |
| v4FP-seq02-1-056 | GNCT | pSEVA29x_lacI_pT7lacO_GNCT | fDG197, fDG140, fDG138, fDG137, fDG142, fDG163, fDG164 |
| v4FP-rand-043 | NTCG | pSEVA29x_lacI_pT7lacO_NTCG | fDG197, fDG138, fDG142, fDG137, fDG140, fDG163, fDG164 |
| v4FP-025-012 | CGTB | pSEVA29x_lacI_pT7lacO_CGTB | fDG197, fDG137, fDG140, fDG142, fDG143, fDG163, fDG164 |
| v4FP-rand-087 | BGNC | pSEVA29x_lacI_pT7lacO_BGNC | fDG197, fDG143, fDG140, fDG138, fDG137, fDG163, fDG164 |
| v4FP-seq02-1-060 | GNTB | pSEVA29x_lacI_pT7lacO_GNTB | fDG197, fDG140, fDG138, fDG142, fDG143, fDG163, fDG164 |
| v4FP-rand-044 | NTCB | pSEVA29x_lacI_pT7lacO_NTCB | fDG197, fDG138, fDG142, fDG137, fDG143, fDG163, fDG164 |
| v4FP-025-016 | CBGT | pSEVA29x_lacI_pT7lacO_CBGT | fDG197, fDG137, fDG143, fDG140, fDG142, fDG163, fDG164 |
| v4FP-seq02-1-107 | TNBC | pSEVA29x_lacI_pT7lacO_TNBC | fDG197, fDG142, fDG138, fDG143, fDG137, fDG163, fDG164 |
| v4FP-rand-050 | GCNT | pSEVA29x_lacI_pT7lacO_GCNT | fDG197, fDG140, fDG137, fDG138, fDG142, fDG163, fDG164 |
| v4FP-rand-118 | TBNG | pSEVA29x_lacI_pT7lacO_TBNG | fDG197, fDG142, fDG143, fDG138, fDG140, fDG163, fDG164 |
| v5FP-rand-078 | BCTGN | pSEVA29x_lacI_pT7lacO_BCTGN | fDG197, fDG143, fDG137, fDG142, fDG140, fDG138, fDG163, fDG164 |
| v4FP-rand-058 | GNBT | pSEVA29x_lacI_pT7lacO_GNBT | fDG197, fDG140, fDG138, fDG143, fDG142, fDG163, fDG164 |
| v4FP-025-057 | GNBC | pSEVA29x_lacI_pT7lacO_GNBC | fDG197, fDG140, fDG138, fDG143, fDG137, fDG163, fDG164 |
| v5o7-rand-1435 | GXTND | pSEVA29x_lacI_pT7lacO_GXTND | fDG197, fDG140, fDG201, fDG142, fDG138, fDG202, fDG163, fDG164 |
| vDG-RF-011c2 | BNU | pSEVA29x_lacI_pT7lacO_BNU | fDG197, fDG143, fDG138, fDG212, fDG163, fDG164 |
| vDG-RF-005c5 | APG | pSEVA29x_lacI_pT7lacO_APG | fDG197, fDG210, fDG211, fDG140, fDG163, fDG164 |
| vDG-RF-029c4 | CNX | pSEVA29x_lacI_pT7lacO_CNX | fDG197, fDG137, fDG138, fDG201, fDG163, fDG164 |
| vDG-RF-053c5 | NTBX | pSEVA29x_lacI_pT7lacO_NTBX | fDG197, fDG138, fDG142, fDG143, fDG201, fDG163, fDG164 |
| vDG-RF-055c7 | PBGXT | pSEVA29x_lacI_pT7lacO_PBGXT | fDG197, fDG211, fDG143, fDG140, fDG201, fDG142, fDG163, fDG164 |
| vDG-RF-058c4 | PCBG | pSEVA29x_lacI_pT7lacO_PCBG | fDG197, fDG211, fDG137, fDG143, fDG140, fDG163, fDG164 |

**Table S5. List of features used for the random forest models.** The features are grouped according to operon context: gene of interest (goi), upstream context, downstream context, and global operon properties. Definitions: (i) the upstream junction is the sequence containing the last 50 nt of upstream gene and the first 50 nt of the goi (first 100 nt of the goi if the goi is the first gene), (ii) the downstream junction is the sequence containing the last 50 nt of the goi and the first 50 nt of the downstream gene (last 100 nt of the goi if the goi is the last gene).

| Operon level | Feature | Description |
|---|---|---|
| Gene of interest | distancePromoter | distance of transcription start site to start codon of coding sequence in base pairs |
| | mRBS | translation initiation rate of monocistronic reference construct |
| | nREMotif | number of RNase E motifs (RNWTT) within the gene |
| Upstream | usRBS | translation initiation rate of monocistronic reference construct of preceding gene |
| | usJunctionFE | minimum free energy of secondary structure for the upstream junction |
| | usJunctionPairedML | mean length of consecutive paired nucleotides in minimum free energy prediction for the upstream junction |
| | usJunctionUnpairedML | mean length of consecutive unpaired nucleotides in minimum free energy prediction for the upstream junction |
| | usREMotif | number of RNase E motifs (RNWTT) within the upstream gene |
| Downstream | dsJunctionFE | minimum free energy of secondary structure for the downstream junction |
| | dsJunctionPairedML | mean length of consecutive paired nucleotides in minimum free energy prediction for the downstream junction |
| | dsJunctionUnpairedML | mean length of consecutive unpaired nucleotides in minimum free energy prediction for the downstream junction |
| | dsREMotif | number of RNase E motifs (RNWTT) within the downstream gene |
| Global | induction | concentration of IPTG used for induction in μM |
| | operonLengthBP | operon length in base pairs from transcription start site last stop codon |
| | overallGC | overall operon GC content of operon |
| | mMPairedL | grand mean of lengths of consecutive paired nucleotides in each gene's minimum free energy RNA structure prediction |
| | mMUnpairedL | grand mean of lengths of consecutive unpaired nucleotides in each gene's minimum free energy RNA structure prediction |
| | gmRelFE | geometric mean of absolute values of all minimum free energies of each gene's secondary RNA structure normalized to gene length for each gene |
| | gmRBS | geometric mean of all translation initiation rates of monocistronic reference construct genes |
